# Genomic patterns of transcription-replication interactions in mouse primary B cells

**DOI:** 10.1101/2021.04.13.439211

**Authors:** Commodore P. St Germain, Hongchang Zhao, Vrishti Sinha, Lionel A. Sanz, Frédéric Chédin, Jacqueline H. Barlow

## Abstract

Conflicts between transcription and replication machinery are a potent source of replication stress and genome stability; however, no technique currently exists to identify endogenous genomic locations prone to transcription-replication interactions. Here, we report a novel method to identify genomic loci prone to transcription-replication interactions termed transcription-replication immunoprecipitation on nascent DNA sequencing, TRIPn-Seq. TRIPn-Seq employs the sequential immunoprecipitation of RNA polymerase 2 phosphorylated at serine 5 (RNAP2s5) followed by enrichment of nascent DNA previously labeled with bromodeoxyuridine. Using TRIPn-Seq, we mapped 1,009 unique transcription-replication interactions (TRIs) in mouse primary B cells characterized by a bimodal pattern of RNAP2s5, bidirectional transcription, an enrichment of RNA:DNA hybrids, and a high probability of forming G-quadruplexes. While TRIs themselves map to early replicating regions, they exhibit enhanced Replication Protein A association and replication fork termination, marks of replication stress. TRIs colocalize with double-strand DNA breaks, are enriched for deletions, and accumulate mutations in tumors. We propose that replication stress at TRIs induces mutations potentially contributing to age-related disease, as well as tumor formation and development.

## INTRODUCTION

In mammals, the DNA replication machinery must accurately duplicate over 6 billion base pairs per cell division. Replication forks navigate through DNA secondary structures, displace and relocate chromatin-associated proteins, and cope with torsional stress all while maintaining processivity and fidelity. Barriers preventing efficient fork progression—termed replication stress—are a potent source of genome instability, a hallmark of cancer (1). A substantial cause of replication stress is transcription, as RNA polymerase (RNAP) must also navigate the same chromatin template while processively transcribing molecules sometimes over 2 Mb long (2–5).

RNA polymerase II (RNAP2) is a large, multi-subunit complex that transcribes all protein coding genes. RNAP2 activity is tightly regulated at individual genes with each stage of transcription—initiation, elongation and termination—coordinated by the recruitment of multiple complexes through the phosphorylation of the carboxy-terminal domain (CTD) repeats. A series of specific and general transcription factors recruit the unphosphorylated RNAP2 to core promoters, a form poised for activity. The cyclin-dependent kinase Cdk7 then phosphorylates RNAP2 at serine 5 (RNAP2s5), allowing the complex to leave the initiation site (6). RNAP2s5 is found in both promoter-proximal paused and elongating complexes throughout the gene body, and interacts with the spliceosome during co-transcriptional splicing (7,8).

Transcription-replication interactions (TRIs) can arise anywhere transcription and replication machineries occur simultaneously on the same DNA template (4). Transcription and replication create single stranded DNA (ssDNA) during template unwinding potentially allowing stable secondary structures such as G-quadruplexes or stem-loops to form in repetitive DNA sequences. Active transcription also stimulates the formation of co-transcriptionally formed RNA:DNA hybrids (R-loops), three-stranded nucleotide structures formed when nascent RNA re-anneals with the DNA template strand behind elongating RNAP2 (9). R-loops and DNA secondary structures can induce replication-associated genome instability, causing fork stalling or collapse (10–12). Stalled forks can potentially resume replication; however, collapsed forks require resolution through various DNA repair pathways (13).

To date, much of the work directly analyzing TRIs has been conducted in bacteria, yeast and *in vitro* (14–16). Studies in bacterial systems show that both co-directional and head-on transcription can induce replication fork stalling and collapse, the latter causing more profound consequences (17–24). TRIs can induce mutations in the form of single basepair substitutions (SBS) as well as short insertions and deletions (indels) at promoters and in protein-coding regions (25).Higher expression and longer gene length correlated with increased accumulation of mutations (28). Both codirectional and convergent TRIs experienced damage in the form of indels and base substitutions throughout the gene, but were focused in the promoter region (25). RNAP elongation can also pause at sites of DNA damage and at regulatory sequences causing it to backtrack. GreA and GreB promote transcript cleavage to release paused and backtracked RNA polymerase in bacteria, and their loss increases replication fork stalling, genome instability and cell death (26).

Studies in mammalian cells have helped elucidate the role that TRIs play in the context of disease. In an episomal system, head-on and codirectional gene transcription both increase plasmid instability and DNA damage in human cell lines (27). Persistent RNAP2 backtracking also induces genome instability in human cells. A recent report found that U2OS and HEK293T cell lines expressing a mutant form of the transcription elongation factor TFIIS that blocks the rescue of RNAP2 “trapped” in a backtracked or paused state—exhibit increased RNAP2 pausing and DNA breaks (28). In transformed and immortalized lymphocytes, transcriptional activity also correlates with aphidicolin-induced genome instability at very long, late-replicating genes located in common fragile sites (3).

However, virtually all studies of TRIs in mammalian cells have employed artificial DNA constructs or chemical agents, and use immortalized cell lines with abnormal genotypes and dysregulated cellular processes allowing for indefinite cellular division (27,29–31). Though valuable insights have been gained using these systems, the location of endogenous TRIs and their impact on genome stability remains largely unexplored in primary mammalian cells. In this study, we use mouse wild-type splenic B cells as a model to study TRIs in primary cells, as naïve B lymphocytes induce a highly regulated wave of transcription and rapid proliferation that can be reliably mimicked *ex vivo* by antigen/cytokine stimulation (32–34).

To define where TRIs naturally arise, we developed Transcription-Replication IP on Nascent DNA coupled with high-throughput sequencing (TRIPn-Seq), a sequential immunoprecipitation (IP) of RNA polymerase II phosphorylated at serine 5 followed by an IP of bromodeoxyuridine (BrdU) labeled nascent DNA. TRIPn-Seq defined the locations of 1,009 independent TRI loci in primary mouse B cells which predominantly occur in early replicating regions. TRIs are characterized by high levels of bidirectional transcription, RNA:DNA hybrid formation, and are strongly enriched for genetic sequences prone to forming secondary structures. We propose that TRIs represent genomic loci enriched for multiple structures hindering replication fork progression, and these persistent obstructions to replication fork progression leads to fork collapse, DNA break formation, and genome instability.

## MATERIALS AND METHODS

### Mice and B cell harvesting

Mouse spleens were isolated from wild-type (WT) mice with a C57BL/6 background. A 70 μm filter was set in a 5 cm petri dish containing 5 mL of cold wash buffer (see all reagent specifics in reagents table below) and the spleens were placed in the filter and gently pushed through with a 5 mL syringe plunger. The cell suspension was transferred to a 50 mL conical tube, an additional 5 mL of wash buffer was added to the petri dish and the remaining material was gently pushed through the filter and transferred to the 50 mL conical. A final 5 mL of wash buffer was used to rinse the petri dish and transfer the remaining cells to the 50 mL conical tube. The 15 mL cell suspension was centrifuged (Sorvall Legend XTR) at 500 g for 5 min and the supernatant was aspirated. The pellet was resuspended in 4 mL ACK lysis buffer by forcefully pipetting 5 times with a 5 mL pipettor and incubating at room temperature (RT) for 4 min and neutralized with 11 mL of wash buffer. Any large visible non-soluble material was carefully removed with a pipette tip. This was centrifuged at 500 g for 5 min. The supernatant was aspirated, and the pellet was resuspended in 925 μL of wash buffer and additional non-soluble material was removed.

To isolate B cells, 75 μL of mouse CD43 Dynabeads per spleen was aliquoted to a FACS tube and combined with 1 mL of wash buffer. This was placed on a magnet for 3 min and the solution was removed and the beads were resuspended in 75 μL of wash buffer per spleen. The 925 μL of cell suspension was added to the beads, capped, and placed on a rotator for 20 min at RT. After incubation, an additional 2 mL of wash buffer was added to the tube, and was pulse centrifuged and placed on a magnet for 5 min. The supernatant was transferred to a new 15 mL conical tube and brought to a total volume of 5-10 mL with wash buffer and cells were counted using a Bio-Rad Tc10.

### B cell culture

B cells were added to pre-warmed B cell media (see reagent list below) and stimulated to proliferate using final concentrations of 5 μg/mL LPS, 2.5 ng/mL IL-4, and 250 ng/mL anti-CD180. Cells were plated in 5 mL volumes at 150,000 cells/mL in 6 well plates and incubated at 37 °C. At 70.5 hours, cells were resuspended with a 5 mL pipettor to dislodge the B cells from the plate and transferred to a 50 mL conical. An aliquot of cell suspension was taken for counting and samples were placed back into the incubator for 20 min with the cap loose back to 37 °C. For BrdU-positive samples, BrdU was added to a final concentration of 10 μM, inverted 4-6 times to mix, then placed back into the incubator with the cap loose for 30 min.

### Cell fixation chromatin isolation

Cells were removed from the incubator and immediately crosslinked with 0.75 % formaldehyde for 2 min at RT, inverting to mix. Cells were quenched with freshly made 1.25 M glycine stock, bringing the solution to 0.125 M glycine, inverting 4-6 times. Cells were washed by immediately centrifuging at 500 g for 5 min, aspirating the supernatant, and resuspending in 50 mL RT PBS. The samples were washed three times and during the third PBS resuspension, the samples were transferred to different conical tubes with the amount of cells that will be used for the ChIP experiment. The samples were again centrifuged at 500 g for 5 min, supernatant was aspirated, and cell pellet was snap frozen in a 100 % EtOH/dry ice bath and stored at −80°C for less than 6 weeks.

### TRIPn and RNAP2s5-ChIP Method 1 (replicates 1 and 2)

Each replicate consists of 2 samples, including a BrdU negative (BrdU-) and a BrdU-positive (BrdU+) sample. This protocol is written for each sample. First, 100 μL of protein G beads were washed with 1 mL TBST, placed on a magnet for 3 min, the supernatant was aspirated, and beads were resuspended in 500 μL TBST. Next, 6 μg of RNAP2s5 antibody was added and placed on a rotator for 4-5 hours at 4°C to be used later.

100-140 million cells were thawed at RT for 10 min, centrifuged at (Sorvall Legend XTR) 500 g for 5 min and supernatant was throrougly aspirated. The cell pellet was resuspended in lysis buffer at 100 μL per 7.5 million cells and vortexed to mix. The lysate was transferred to the sonication tubes at 200-300 μL per tube and sonicated on low for 3 cycles at 30 secs on, 30 secs off at 4°C. This produced a smear of DNA products of sizes primarily ranging from 500-2,000 bp as visualized on an 1.25 % TAE agarose gel. The sonicated samples were centrifuged (Eppendorf 5424R) at 13,000 g for 5 min and the supernatant was transferred to a conical tube. For gel samples to visualize fragment lengths, 1 μL gel of sonicated cell solution was added to 19 μL PBS and proteinase K was added to a final concentration of 100 μg/mL and placed in a shaking dry bath at 55°C, for 14 hours, at 500 rpm.

The supernatant was diluted 1:10 with TBST, mixed, and portions were transferred to an Amicon concentrator tube. This was centrifuged (Sorvall Legend XTR) at 4,000 g until the remaining solution in the top chamber was ~1 mL. This was pipetted to mix and dislodge any material that was bound to the filter surface and additional sonicated chromatin solution was added and centrifuged again. This was repeated until the total amount was reduced to ~1.5 mL.

The protein G / RNAP2s5 antibody mixture was pulse centrifuged, placed on a magnet for 5 min and the supernatant was aspirated and discarded. The solution in the Amicon filter was pipetted again to mix and dislodge any material bound to the filter and transferred to the 1.5 mL tube that contained the washed protein G / antibody combination. This was placed on a rotator at 4 °C O/N.

The next day the samples were pulse centrifuged, placed on a magnet for 5 min and the supernatant was aspirated and discarded. The beads were gently resuspended with a 1,000 μL micropipette with 1 mL of low salt buffer, transferred to a new 1.5 mL tube, and placed on a magnet for 5 min. This step was repeated with high salt buffer and then LiCl buffer also transferring to new tubes after each resuspension. The LiCl buffer was discarded and beads were resuspended with 60 μL of elution buffer and placed on a shaking dry bath at 45 °C for 20 min at 700 rpm. This was placed on a magnet for 3 min and the supernatant was transferred to a new tube. The beads were rinsed with an additional 60 μL of elution buffer, placed on a magnet for 3 min, and the additional supernatant was added to the previous 60 μL of elution. This was stored this in 4 °C until ready for overnight reverse crosslinking. To reverse crosslink, 280 μL of TBST and 100 μg/mL of proteinase K was added and placed on shaking dry bath at 55 °C for 14 hours at 500 rpm.

**[Specifically for TRIPn, not for RNAP2s5-ChIP:** 20 μL of protein G beads were washed with 500 μL TBST, placed on a magnet for 3 min, the supernatant was aspirated, and beads were resuspended in 100 uL TBST. 1 μg of anti-BrdU antibody was added and placed on a rotator for 4-5 hours at 4 °C to be used later.]

The next day the samples were pulse centrifuged, transferred to sonication tubes and sonicated on high for 14 cycles at 15 secs on, 45 secs off at 4 °C to reduce the fragment size to 200-500 bp. The samples were pulse centrifuged and transferred to a new 1.5 mL tube. 400 μL of phenol/chloroform was added and vortexed on high for 10 secs. Samples were pulse centrifuged, transferred to a phase lock tube, and centrifuged again for 5 min at 13,000 g. The top layer was transferred to a new 1.5 mL tube, 1,128 of 100 % EtOH, 41 uL of 3M sodium acetate, and 2 μL of glycogen were added and vortexed on high for 10 secs. This was incubated at −80 °C for 1-2 hours, centrifuged (Eppendorf 5424R) for 30 min at 13,000 g at 4 °C, the supernatant was discarded, 500 μL of EtOH was added without dissolving the pellet, and centrifuged (Eppendorf 5424R) again for 15 min at 13,000 g at 4 °C. The supernatant was discarded, and the pellet was dried until all the EtOH was evaporated—about 15 min and no more than 20 min—by placing the 1.5 mL tube upside down at a 45° degree angle, ensuring the pellet does not slide down the tube. The pellet was resuspended in 52 μL of 0.1X TE for 30 min while briefly vortexing and centrifuging (Eppendorf 5424R) every 10 min. The concentration was measured by Qubit.

The NEBNext End Prep protocol and reagents were used followed by adapter ligation. The adapters were diluted 1:10 in 10 mM Tris-HCl pH 8.0 and 10 mM NaCl. The NEBNext Ultra II DNA Library Prep Kit for Illumina directions were used for an AmpureXP clean-up starting with 86 μL (92 % of total volume) AmpureXP beads.

**[Specifically for TRIPn, not for RNAP2s5-ChIP:** The DNA attached to the AmpureXP beads was eluted with 30 μL of water and transferred to a 1.5 mL tube. 120 μL of TBST was added to the tube, heated in a shaking dry bath at 95°C for 5 min and immediately placed on ice.

The protein G / BrdU antibody mixture was pulse centrifuged, placed on a magnet for 5 min and the supernatant was aspirated and discarded. The solution from the AmpureXP clean-up was transferred to the tube that contained the washed protein G / BrdU antibody combination. This was placed on a rotator at 4°C O/N.

The next day the samples were pulse centrifuged, placed on a magnet for 5 min and the supernatant was aspirated and discarded. The beads were gently resuspended with a 1,000 μL micropipette with 500 μL low salt buffer, transferred to a new 1.5 mL tube, and placed on a magnet for 5 min. This step was repeated with high salt buffer and then LiCl buffer also transferring to new tubes after each resuspension. The LiCl buffer was discarded and the beads were resuspended with 25 μL of elution buffer and place on a shaking dry bath at 45 °C for 20 min at 700 rpm. This was placed on a magnet for 3 min and the supernatant was transferred to a new tube. The beads were rinsed with an additional 25 μL of elution buffer, placed on a magnet for 3 min, and additional supernatant was added to the previous 25 μL of elution.

50 μL of water was added and a two-step AmpureXP clean-up was performed a to remove large and small fragments. 55 μL beads (55 % of total volume) was first added, mixed, and incubated for 5 min, and placed on a magnet for 5 min. The supernatant was collected and an additional 25 μL beads was added (80 % total Ampure solution) and the rest of the procedure was performed as indicated in the NEB protocol.]

The DNA bound to the AmpureXP beads was eluted in 32 μL of 0.1X TE. 1 μL was used to perform a test qPCR amplification to ensure an adequate amount of material in the library amplification. 15 μL was stored at −20 °C and 15 μL was used for library amplification (this can be stored at −20 °C for future amplification). Library amplification was also performed using the NEBNext Ultra II DNA Library Prep Kit for Illumina using NEB Illumina Adaptors and was purified using the Ampure XP beads. 50 μL of water was added to the 50 μL of PCR product and a two-step AmpureXP clean-up was performed the same as above using 58 μL of beads for the first step and 20 μL of beads for the second step and eluted in 17 μL of 0.1X TE. One μL was used for a Qubit measurement for DNA concentration and 1 μL was used for a Bioanalyzer analysis for fragment size. The library was submitted to Novogene for sequencing on an Illumina Hiseq Platform PE150.

### TRIPn and RNAP2s5-ChIP Method 2 (replicate 3)

Each replicate consists of 2 samples, 1 BrdU- and 1 BrdU+ and this protocol is written for 1 sample. 10 million cells were thawed at RT for 10 min, centrifuged at 500 g for 5 min and any small amount of fluid was aspirated. The cell pellet was resuspended in RIPA+ buffer at 300 μL per 10 million cells, vortexed to mix, and transferred to the sonication tubes at 200-300 μL per tube. Samples were sonicated on high at 4 °C for 8 cycles at 15 secs on, 45 secs off, rotating sonicator positions after 4 cycles. To confirm sonication efficiency, 2 μL gel of sonicated cell solution was added to 18 μL PBS and proteinase K (final concentration 100 μg/mL) and placed in a shaking dry bath at 55 °C, for 14 hours, at 500 rpm. Successful samples produced a smear of DNA products on an 1.25 % TAE agarose gel, concentrated between 200-1,500 bp. Sonicated samples were centrifuged at 13,000 g for 5 min and the supernatant was transferred to a 1.5 mL tube. The supernatant was diluted 1:5 with TBST and 1 μg of RNAP2s5 antibody was added per 275 of diluted chromatin supernatant. This was placed on a rotator at 4 °C O/N.

10 % of the IP volume of protein G beads were washed with 1,000 μL TBST, placed on a magnet for 3 min and the supernatant was discarded. The RNAP2s5 IP solution from the previous night was pulse centrifuged and pipetted into the protein G beads and pipetted to mix. This was placed on a rotator for 4-5 hours at 4 °C.

Samples were pulse centrifuged, placed on a magnet for 5 min and the supernatant was aspirated and discarded. Beads were gently resuspended with a 1,000 μL micropipette with 1 mL of low salt buffer, transferred to a new 1.5 mL tube, and placed on a magnet for 5 min. This step was repeated with high salt buffer and then LiCl buffer also transferring to new tubes after each resuspension. The LiCl buffer was discarded and beads were resuspended with 100 uL of elution buffer and place on a shaking dry bath at 45 °C for 20 min at 700 rpm. The beads were placed on a magnet for 3 min and supernatant was transferred to a new tube. Beads were washed with an additional 100 μL of elution buffer, placed on a magnet for 3 min, and the supernatant was added to the previous 100 μL of elution. This was stored in 4 °C until ready for overnight reverse crosslinking. To reverse crosslink, 200 μL of PBS and 100 μg/mL of proteinase K was added and placed on shaking dry bath at 55 °C for 14 hours at 500 rpm.

The next day the samples were pulse centrifuged, transferred to sonication tubes and sonicated on high for 10 cycles at 15 secs on, 45 secs off at 4 °C to reduce the fragment size to 200-500 bp. The samples were pulse centrifuged and transferred to a new 1.5 mL tube. 400 μL of phenol/chloroform was added and vortexed on high for 10 secs. This was pulse centrifuged, transferred to a phase lock tube, and centrifuged again for 5 min at 13,000 g. The top layer was transferred to a new 1.5 mL tube, 1128 of 100 % EtOH, 41 μL of 3M sodium acetate, and 2 uL of glycogen was added and vortexed on high for 10 secs. This was incubated at −80 °C for 1-2 hours, centrifuged (Eppendorf 5424R) for 30 min at 13,000 g at 4 °C, the supernatant was discarded, 500 mL of EtOH was added without dissolving the pellet, and centrifuged (Eppendorf 5424R) again for 15 min at 13,000 g at 4 °C. The supernatant was discarded, and the pellet was dried until all the EtOH was evaporated for 15 but no more than 20 min by placing the 1.5 mL tube upside down at a ~45° degree angle ensuring the pellet doesn’t slide down the tube. The pellet was resuspended in 52 μL of 0.1X TE for 30 min while briefly vortexing and centrifuging every 10 min. The concentration was measured by Qubit.

The NEBNext End Prep protocol and reagents were used followed by adapter ligation. Adapters were diluted 1:10 in 10 mM Tris-HCl pH 8.0 and 10 mM NaCl. The NEBNext Ultra II DNA Library Prep Kit for Illumina directions were used for an AmpureXP clean-up starting with 86 μL (92 % of total volume) AmpureXP beads.

#### [Specifically for TRIPn, not for RNAP2s5-ChIP

The DNA attached to the AmpureXP beads was eluted with 50 μL of 0.1X TE and transferred to a 1.5 mL tube. 150 μL of TBST was added to the tube, heated in a shaking dry bath at 95°C for 5 min and immediately placed on ice. 1 μg of BrdU antibody was added and placed on a rotator at 4°C overnight.

The next day 20 μL of protein G beads were washed with 500 μL TBST, placed on a magnet for 3 min, supernatant was discarded, and beads were resuspended in 100 μL TBST. The BrdU IP solution from the previous night was pulse centrifuged and pipetted into the protein G beads and pipetted to mix. The protein G-BrdU IP mixture was placed on a rotator for 4-5 hours at 4°C.

After incubation, the samples were pulse centrifuged, placed on a magnet for 5 min and the supernatant was aspirated and discarded. The beads were gently resuspended with a 1,000 μL micropipette with 500 μL low salt buffer, transferred to a new 1.5 mL tube, and placed on a magnet for 5 min. This step was repeated with high salt buffer and then LiCl buffer also transferring to new tubes after each resuspension. We discarded the LiCl buffer and resuspended with 25 μL of elution buffer and place on a shaking dry bath at 45°C for 20 min at 700 rpm. This was placed on a magnet for 3 min and the supernatant was transferred to a new tube. The beads were rinsed with an additional 25 μL of elution buffer, placed on a magnet for 3 min, and the additional supernatant was added to the previous 25 μL of elution.

100 μL of water was added and a two-step AmpureXP clean-up was performed to remove large and small fragments. 82.5 μL beads (55 % of total volume) was first added, mixed, and incubated for 5 min, and placed on a magnet for 5 min. The supernatant was collected and saved and an additional 37.5 μL beads (80 % total Ampure solution) was added, and the rest of the procedure was performed as indicated in the NEB protocol.]

The DNA bound to the AmpureXP beads was eluted in 32 μL of 0.1X TE. One μL was used to perform a test qPCR amplification to ensure an adequate amount of material in the library amplification. Fifteen μL was stored in −20°C and 15 μL was used for library amplification (this can be stored at −20°C for future amplification). Library amplification was also performed using the NEBNext Ultra II DNA Library Prep Kit for Illumina using NEB Illumina Adaptors and was purified using the Ampure XP beads. Fifty μL of water was added to the 50 μL of PCR product and a two-step AmpureXP clean-up was performed the same as above using 58 μL of beads for the first step and 20 μL of beads for the second step and eluted in 17 μL of 0.1X TE. One μL was used for a Qubit measurement for DNA concentration and 1 μL was used for a Bioanalyzer analysis for fragment size. The library was submitted to Novogene for sequencing on an Illumina Hiseq Platform PE150.

### Flow Cytometry - Replication Timing for S Phase Cells

B cells were isolated and grown as described above. For the control sample, HU was added to a final concentration of 10 μM. At the indicated time points, 2 mL of cells were aliquoted to a 15 mL conical and centrifuged (Eppendorf 5424R) at 500 g for 5 min, the supernatant was aspirated, and the cells were resuspended in 1.5 mL of cold PBS. The cells were permeabilized and fixed by adding 3.5 mL of 4 °C, 100 % EtOH slowly while mixing and incubating at −20°C for 20 min. The cells were washed twice after centrifugation at 850 g for 5 min, discarding the supernatant, and resuspending in 5 mL of PBS. After the second wash, the cells were resuspended in 2 mL of PBS and propidium iodide (PI) was added to a final concentration of 10 μg/mL. PI incorporation was measured on a BD Canto II, comparing experimental cells to HU treated control cells.

### DNA-RNA Immunoprecipitation (DRIP)

DRIP-Seq was performed as described previously (35).

## REAGENTS

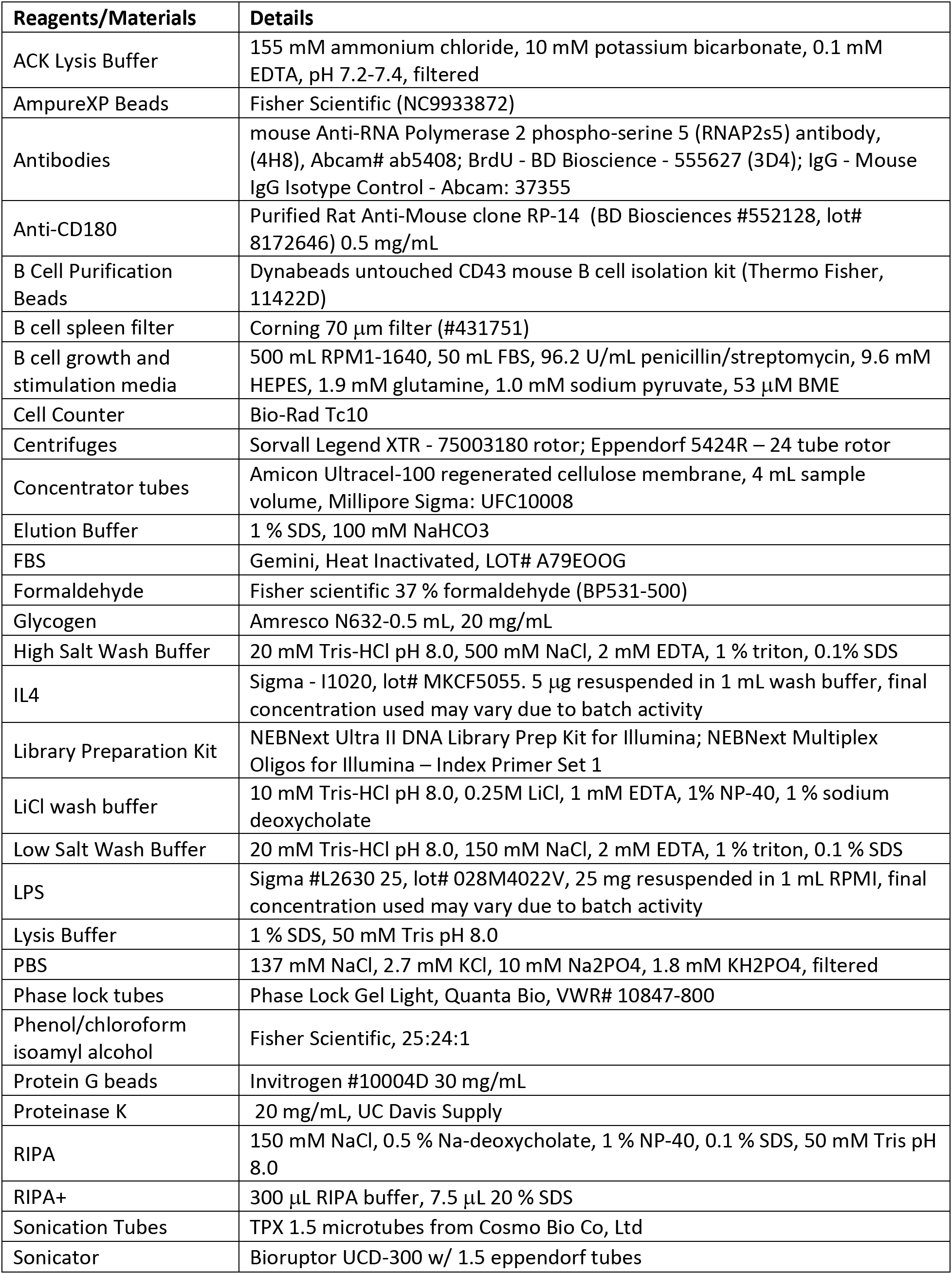
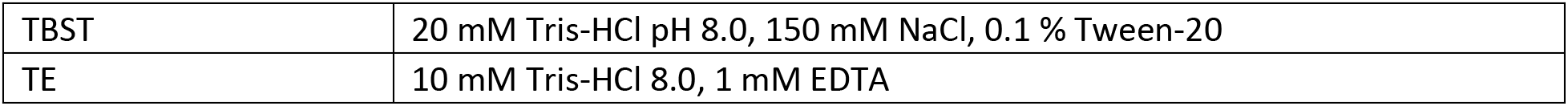

## COMPUTATIONAL METHODS AND RESOURCES

### Programs to perform general computational and transformation functions

Bedtools v 2.26.0 (https://bedtools.readthedocs.io/en/latest/) (36); Python v 2.7.15rc1 (https://www.python.org/) (37); Python v 3.6.9 (https://www.python.org/) (38); SciPy v 1.0: matplotlib, pandas, numpy (https://www.scipy.org) (39–43); UCSC table browser (44); UCSC executable programs: wigToBigWig, liftOver, bigWigToBedGraph (http://hgdownload.soe.ucsc.edu/admin/exe/) (45)

### Basic fastq to bigwig workflow

If using downloaded SRA data, fastq-dump 2.7.0 (https://ncbi.github.io/sra-tools/fastq-dump.html) was used to generate the fastq from SRA file (46). Fastq reads were trimmed with Trimmomatic v 0.36 (http://www.usadellab.org/cms/?page=trimmomatic) (47) and aligned to the mm10 mouse genome (48) with Bowtie2 v 2.2.8 (http://bowtie-bio.sourceforge.net/bowtie2/index.shtml) (49). Unmapped reads, unpaired reads, and PCR duplicates were discarded, and the file was converted into BAM format using Samtools v 1.9 (http://www.htslib.org/) (50). The BAM file was converted to a bedgraph file using bedtools genomecov and normalized by reads per million. The bedgraph file was converted to a bigwig file and visualized using the UCSC genome browser (https://genome.ucsc.edu/) (51).

### TRI Identification

Peaks were called on three independent biological replicate experimental (BrdU+) and three control (BrdU-) TRIPn-Seq BAM files using macs2 callpeak v 2.2.5 (https://github.com/macs3-project/MACS) (52) using the parameters: -f BAMPE –nomodel -g mm. The experimental and control counts were then analyzed and compared to each other using R 4.0.2 (https://www.r-project.org/) (53) and DiffBind v 2.16.0 (https://bioconductor.org/packages/release/bioc/html/DiffBind.html) (54,55) using default parameters except: dba.count(OBJECT, minOverlap=0). Additional TSS peaks were generated by creating 1kb windows +/− 1kb from the TSSs in sliding increments of 100 bp and each peak set was individually analyzed with the edgeR analysis of DiffBind. All peaks that were measured to have differential signal between experimental and control with an FDR ≤ 0.05 were combined and merged and the highest FDR was kept.

### Overlap/Association Experiments

Genomic Association Tester v 1.0 (GAT) (https://gat.readthedocs.io/en/latest/) (56) was used to test association between TRIs and TSSs (57) in 1 kb windows, 24 hr NT END-seq peaks (58), and CpG islands (59,60). The workspace was restricted to the feature (TSSs, END-seq peaks, CpG) merged with all genes that overlapped with RNAP2s5 ChIP-Seq peaks +500 bp upstream of the TSS. GAT was also used to test the overlap in kbp between TRIs and early replicating fragile sites (ERFSs) restricted to early replicating regions and (61) and common fragile sites (CFSs) (62–67) in bp because of the large size difference.

### TRI TSS Orientations

Divergent promoters were plus and minus strand TRITSSs that were within 3kb of their upstream regions using bedtools intersect. Convergent promoters were plus and minus strand TRITSSs that were within 3kb of their downstream regions using bedtools intersect. Bidirectional transcription with only one annotated MGI gene were TRITSSs in 3kb regions that intersected with GRO-Seq peaks that were on both the plus and minus strands. Peaks were called using MACS2 with the same settings as TRIs.

### Profile Plots and Heatmaps

Profile plots and heatmaps were created by using deepTools 3.1.3 (https://deeptools.readthedocs.io/en/develop/) plotMatrix, plotProfile, and plotHeatmap while removing the gaps and blacklisted regions and plotting the median and standard error. In order to compare the same amount of measurements between the regions of interest, cTSSs were shuffled and a number of regions were plotted to match the same number of TRI or TRITSS regions. (68)

### TimEX

DeepTools bamCompare was used to calculate the log2 difference between the reads in the activated B cell BAM file and the resting B cell BAM file (GSE116318 (69)) with the following parameters: --operation log2 --smoothLength 500 --binSize 100 --bl mm10.GAP.BLACKLIST.BED --effectiveGenomeSize 2652783500.

### OK-Seq

DeepTools bamCoverage was used to count reads in the forward strand BAM file in 1 kb windows using the following parameters: -bs 1000, --bl mm10.GAP.BLACKLIST.BED --effectiveGenomeSize 2652783500 - of bedgraph. The mm10 genome was partitioned into 1 kb windows. The reads from bamCoverage were mapped to the mm10 genome 1kb windows using the mean. This was repeated for the reverse strand BAM file. Using the files from the forward and reverse strands, a new replication fork direction (RFD) file in bedgraph format was created using this calculation (R - F) / (R + F) then converted to a bigwig file (70).

### TRI Overlap with Tumor Mutations

The Mouse Tumor Biology Database (71) provided us with data that listed gene names associated with mutations found in sequenced mouse tumors (23,625 mutations) and unique gene names (6525 genes) were extracted. The 1,198 TRI genes were compared with the tumor mutation unique gene name list. A complete gene list was constructed by taking all MGI annotations (302,974 annotations) and removing annotations that were predicted genes, TSS only, and did not contain precise coordinates or a specific strand and 28,398 genes remained. 1,198 random genes were extracted and compared with the tumor mutation gene list. We performed the random extraction and comparison 999 times. For analysis of mutation type, all mutations were kept including non-unique gene names. The same procedure was performed using data from Sleeping Beauty Cancer Driver Database (72) but using 1,231 identified cancer driver genes.

### Variant Analysis

bcftools v 1.9 (http://samtools.github.io/bcftools/bcftools.html) mpileup was used to create genotype likelihoods on 4 different whole genome sequencing files GSM3227968, GSM3227969 (69), GSM4098725 and GSM4098729 (73) using the parameters: -B -Ou -f mm10.fa --max-depth 4000 --max-idepth 2000. Next bcftools call was used to identify variant sites using the parameters: mA -Ob. The bcf files were then indexed using bcftools index. The regions of interest used for TRITSSs and cTSSs were +/− 500 bp from the annotated TSS from the MGI and merged if containing overlapping regions. The regions of interest used for genes were the entire length of the annotated gene +500 bp upstream of the TSS (74). Next, the 4 bcf files were converted to vcf files. The 5 regions of interest were extracted for each of the 4 vcf files using bcftools convert with the following parameters: --threads 8 -O z -R [regions of interest] for a total of 20 files. The files were sorted using bcftools sort -O z and indexed using bcftools index. The 4 TRI files were then combined using bcftools merge -O v and this was repeated for the TRITSS, cTSS, TRI full gene, control full gene files. Mutational signatures were generated using the merged files using SigProfilerMatrixGenerator v 1.1 (https://github.com/AlexandrovLab/SigProfilerMatrixGenerator) (75). To measure enrichment or depletion of single basepair substitutions and indels at TRIs, cTSSs, and TSSs that had RNAP2s5 signal within genic regions, we counted the variants within 1kb regions centered at the 1,198 TRITSSs and then randomly sampled 1,198 regions of the 12,957 cTSSs and counted the variants and repeated the random sampling for a total of 999 times (76). The same procedure was repeated for TRI genes, cTSS genes, and all genes with RNAP2s5 signal except the samples were split into 1kb windows to normalize for gene length.

### G4 Analysis

G4 quadruplex formation was predicted using G4 Hunter v 3.0 in 400 bp windows using the -w 25 -s 1.4 parameters (https://github.com/AnimaTardeb/G4Hunter) (77).

### GC Skew

GC skew was calculated by measuring (G-C)/(G+C) in 200 bp regions using bedtools nuc using a sliding window of 1 basepair where the result of the calculation corresponds to the center of the 200 bp region (78).

### Motif Extraction

Motifs were extracted using HOMER v 4.9.1 findMotifsGenome.pl in 400 bp windows (http://homer.ucsd.edu/homer/) (79).

### Gene Enrichment Analysis

Gene enrichment analysis was performed using MouseMine (www.mousemine.org) (80), g:Profiler (https://biit.cs.ut.ee/gprofiler/gost) (81), and Enrichr (https://maayanlab.cloud/Enrichr/) (82,83).

## STATISTICAL ANALYSES

Statistical analyses between the profile plot signals was performed similar to the DeepTools method of creating the profile plots. Further normalization was not needed because the comparisons between TRITSSs and cTSSs were from the same already normalized bigwig files. First the bigwigs were converted to bedgraphs, splitting the region of interest (30kb for TimEX, 5kb for GC-Skew, 1kb for RPA ChIP-Seq, GC percent, and DNA methylation) into smaller windows (1kb for TimEX, 10 bp for all others). Next, the the median bedgraph signal intensity was mapped onto the new smaller windows using bedtools map, then the mean signal of values was calculated across the entire region of interest. The difference in the calculated values in the regions of interest between 1,198 TRITSSs and 12,957 cTSSs was expressed using a p-value calculated using Wilcoxon ranksums in SciPy (84,85). The p-values for enrichment and depletion of single basepair substitutions and indels and all overlap analyses was calculated using the empirical p-value where p = (r + 1) / (n + 1) where r is the number of times the hypothesis tested is false and n is the total number of comparisons (56,86). The enrichment or depletion of TRI gene type, MTB mutation type, and COSMIC mutation frequency were calculated using the hypergeometric distribution p-value (87).

## RESULTS

### Transcription-replication immunoprecipitation on nascent DNA followed by whole-genome sequencing (TRIPn-Seq)

To map TRIs genome-wide, we developed TRIPn-Seq, a sequential IP coupled to genome-wide sequencing that first isolates RNAP2s5-bound DNA, then enriches for nascent DNA (nDNA) labeled with BrdU (Figure 1a). We stimulated freshly-isolated mouse splenic B cells for rapid proliferation for 72 hours, then pulse labeled with BrdU. Colocalization of RNAP2s5 and nDNA can occur by replication encountering active transcription complexes; however, RNAP2s5 can also reload onto DNA post-replication. To minimize detection of RNAP2s5-BrdU co-IP from post-replication, we transiently labeled DNA for 30 minutes. Cells were then crosslinked, sonicated to produce chromatin fragments, then subjected to the first IP for RNAP2s5. Transcription and replication utilize very large protein complexes, therefore they may be separated by a significant amount of DNA. Both processes also create supercoiled DNA which may induce even further linear separation on the DNA template. Thus, the distance between transcription and replication markers—RNAP2s5 and BrdU—is likely longer than typical ChIP-Seq experiment chromatin fragments which are between 200-500 bp. We chose an initial chromatin fragment size of 300-1,500 bp to increase isolation of DNA fragments with both nDNA and RNAP2s5, while keeping the fragment size short enough to still retain high spatial resolution. The RNAP2s5-IP chromatin eluted DNA was sonicated again, ligated to sequencing library adapters, and used for the second IP for BrdU. Library preparation was completed on the BrdU-IP eluate and subjected to high-throughput sequencing.

**Figure 1.**
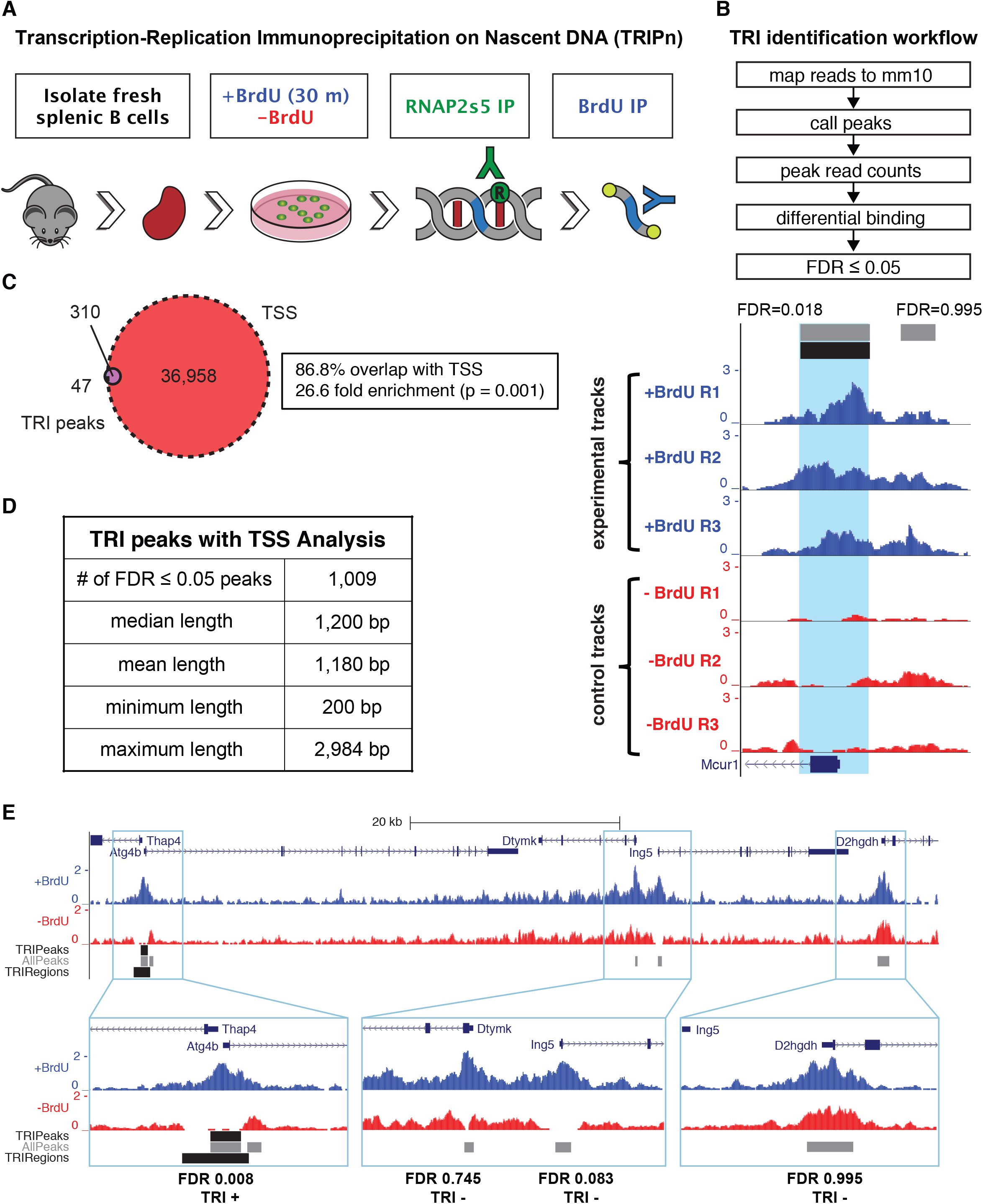
Transcription-Replication ImmunoPrecipitation on Nascent DNA followed by high throughput sequencing (TRIPn-Seq). **(A)** Benchtop workflow of TRIPn-Seq. Mice spleen were extracted, and B cells were isolated and stimulated for growth. Prior to harvesting, BrdU was added to the medium for 30 min. The first IP was for RNAP2s5. Library adapters were added and then the second IP was performed for BrdU. **(B)** Bioinformatic workflow of TRIPn-Seq and representative UCSC genome browser tracks of RPM normalized reads for three TRIPn-Seq experiments (blue) and three controls (red). All experiments are performed in wild-type (WT) mouse B cells (mBCs) unless noted otherwise. Gray bars are areas analyzed along with the calculated FDR by DiffBind and edgeR of differential signal between experimental and control. Black bar and light blue shading are areas that are considered TRIs because the FDR ≤ 0.05. The black bar with arrows at the bottom shows the Mcur1 gene and direction of transcription. **(C)** Venn diagram showing the overlap between TRIs and transcription start sites (TSS), showing empirical p-value. **(D)** Properties of 1,009 TRIs including additional analyzed TSSs. **(E)** Representative UCSC genome browser tracks showing TRI positive regions with FDR ≤ 0.05 and TRI negative regions (FDR > 0.05) and their overlap with TSSs. AllPeaks (gray) are all peaks found using MACS2 and analyzed with DiffBind, TRIPeaks (upper black bar) are peaks with differential signal compared to control with an FDR ≤ 0.05, TRIRegions (lower black bar) also includes analyzed TSSs.

To identify genomic areas specifically enriched for both RNAP2s5 and BrdU, we performed TRIPn-Seq in triplicate on cells incubated with BrdU and compared to those without BrdU. Two different ChIP methods (described as Method 1 and Method 2) were employed for TRIPn-Seq because optimization of TRIPn-Seq revealed that different ChIP conditions gave distinct RNAP2s5 ChIP-Seq peak sets. Representative UCSC genome browser tracks show that Method 1 and Method 2 both exhibited strong RNAP2s5 signal at gene promoters, but Method 2 showed additional transcription-associated peaks not found in Method 1 (Supplementary Figure S1a, yellow). Analysis of the three biological replicates showed BrdU-positive and BrdU-negative samples strongly clustered regardless of ChIP method (Supplementary figure S1b). We mapped the sequencing data from three biological replicates and three controls to the mouse genome assembly mm10 and called peaks. We measured differential binding comparing the peaks pooled from experimental samples against controls and initially found 357 TRI loci with significantly more signal over the control using a false discovery rate (FDR) cutoff of ≤ 0.05 (Figure 1b, Supplementary Figure S1c).

#### TRIPn-Seq signal correlates with active TSSs

Visual inspection of TRI peaks on the UCSC genome browser indicated a strong overlap with transcription start sites (TSSs, Figure 1c). Indeed, 86.8% of TRIs (334/357) overlap with TSSs curated from Mouse Genome Informatics (MGI), a 26.6 fold enrichment over random chance (57). To define additional TRI peaks near TSSs, we analyzed TRIPn-Seq signal +/− 1kb from all TSSs in 800 bp sliding windows with 100 bp steps and each sliding window set was analyzed individually. Loci with FDR ≤ 0.05 were merged with the original 357 TRI loci for a total of 1,009 TRI loci (Figure 1d). We found that TRI experimental peaks with FDR ranging from 0.008 to 0.995 showed similar TRIPn-Seq signals, the difference between low and high FDR was in the difference in signal between the experimental samples and controls (Figure 1e).

To define the number of genes associated with TRIs, we analyzed +/− 1.5 kb flanking the TRI peak centers and found that some TRIs overlap multiple genes, such that the 1,009 TRI loci are associated with 1,221 total TRIs when intersecting nearby annotated genes on the plus and minus strands (1,198 unique genes, Supplementary Table S1). For controls, we used 12,957 control TSSs (cTSSs) which have RNAP2s5 signal but were not considered TRIs (FDR > 0.05; Figure 1c,e). TRIs do not center precisely at TSSs, therefore we also performed all downstream analyses centering the 1,198 TSS-overlapping TRIs on the TSSs (TRITSSs) for more accurate comparison to cTSSs.

### General properties of TRIs and their associated genes

We next compared the length of genes associated with TRIs to all RNAP2s5-bound genes. We found that TRI-associated genes are consistently longer than RNAP2s5-bound genes (median length 41,017 bp for TRIs versus 17,694 bp for cTSS genes; p = 3.45 x 10^−65^). When we restricted analysis to protein coding genes only, the median length of TRI genes was still significantly longer than controls (41,071 bp vs. 20,830 bp; p = 6.06 x 10^−40^, Supplementary Figure S1d). We analyzed the type of genes that TRIs overlapped and found an enrichment of protein-coding genes and a depletion of pseudogenes and long non-coding RNA genes compared to all genes (Supplementary Table S2). We also observed a depletion of micro RNA (miRNA) genes. This may be an underrepresentation as miRNAs are often found in clusters, and our analyses associated a single miRNA per TRI. Additionally, we found that TRI frequency per chromosome strongly correlated with the number of genes per chromosome. However, there was a depletion of TRIs on ChrX and Chr7 (Supplementary Figure S1e).

In addition to identifying unique RNAP2s5 ChIP-Seq peaks, the Method 2 signal pattern was similar to reports of bimodal RNAP2 signals from high-coverage ChIP-Seq experiments and single nucleotide resolution footprinting studies, indicating that Method 2 provides higher resolution for RNAP2s5 signal (88,89). Using the Method 2 RNAP2s5 ChIP-Seq data, we found that cTSSs had a single RNAP2s5 peak in the center while TRIs had two flanking RNAP2s5 signals (Figure 2a, left). TRI peaks were greatly outnumbered by cTSS peaks, potentially confounding signal comparisons along individual loci by metagene analysis. To compare the same number of peaks, cTSSs were randomly shuffled and similar number of regions were plotted for cTSSs and TRIs. Heatmaps showed the majority of TRI/TRITSS loci exhibit bimodal distribution of RNAP2s5 signal, while cTSSs harbored a single central peak (Figure 2a, right). We also measured RNAP2s5 occupancy throughout the entire gene body and at transcription termination sites (TTSs), and found that genes associated with cTSSs had more RNAP2s5 occupancy than TRITSS genes (Supplementary Figure S2a).

**Figure 2.**
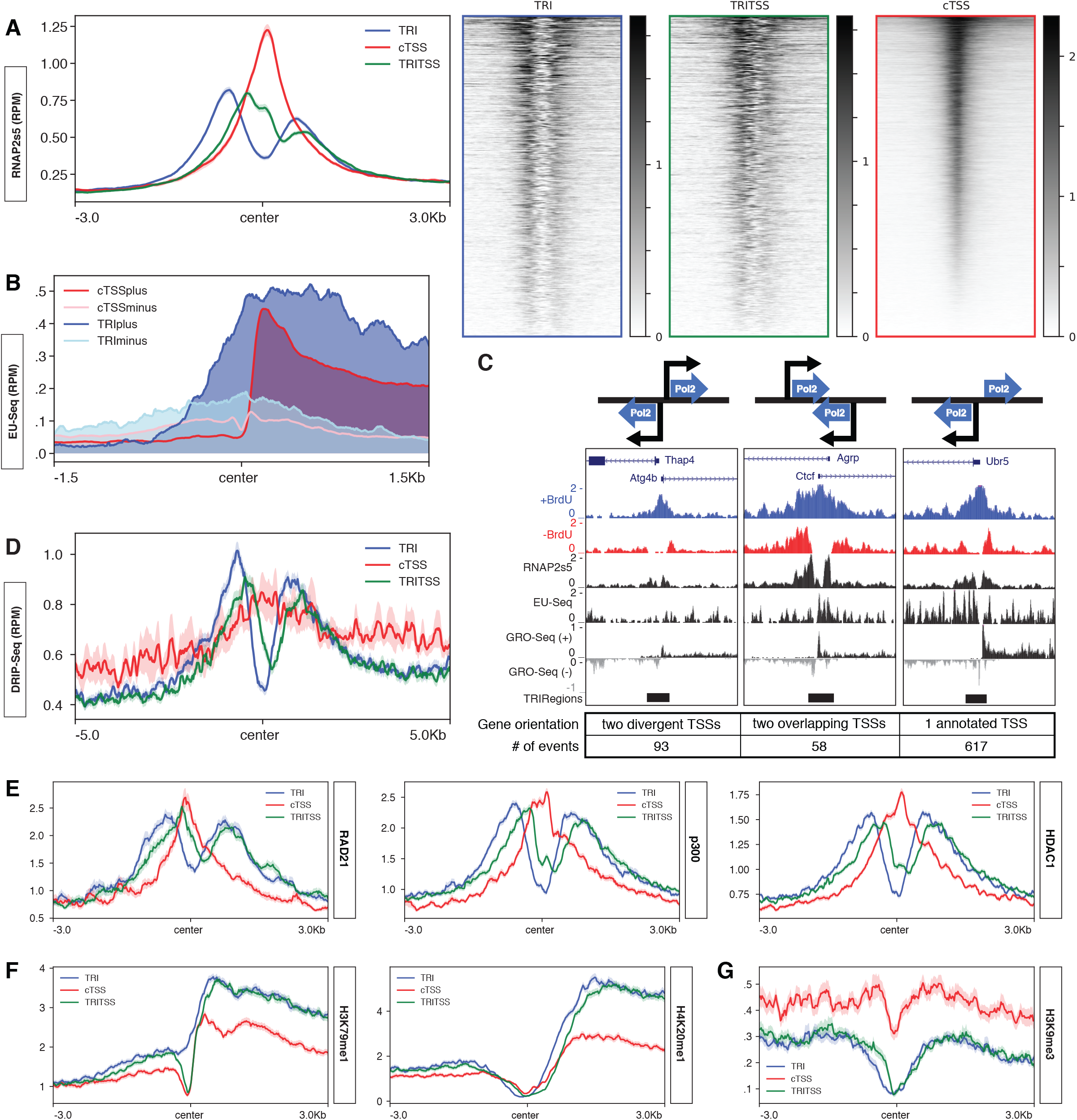
Characterization of transcriptional activity at TRIs. **(A)** Median Method 2 RNAP2s5 ChIP-Seq signal profiles centered near TSSs for TRIs (blue, n=1,221), TRITSSs (green, n=1,198), and cTSSs (red, n=12,957). Median Method 2 RNAP2s5 ChIP-Seq signal heatmaps centered near TSSs for TRIs (blue, n=1,221), TRITSSs (green, n=1,198), and cTSSs (red, n=1,218) **(B)** Median EU-Seq signal profiles centered near TSSs for plus strand nascent RNA at TRIs (blue), minus strand nascent RNA at TRIs (light blue), plus strand nascent RNA at cTSSs (red), and minus strand nascent RNA at cTSSs (pink). **(C)** Graphic representation of possible RNAP2s5 orientations and representative UCSC genome browser tracks of RPM normalized reads of (from top to bottom) Mouse Genome Informatics gene annotation, TRI region, TRI control, RNAP2s5 ChIP-Seq, EU-Seq, and GRO-Seq, and TRI peak for Ctcf, Gm5914, Thap4, Atg4b, and Ubr5. **(D)** Median DRIP-Seq signal profiles centered near TSSs for TRIs, TRITSSs, and cTSSs. **(E)** Median RAD21, P300, and HDAC1 ChIP-Seq signal profiles in RPKM for TRIs, TRITSSs, and cTSSs. **(F)** Median H3K9me3, H3K79me1, and H4K20me1 ChIP-Seq signal profiles in RPKM for TRIs, TRITSSs, and cTSSs. **(G)** Median H3K9me3 ChIP-Seq signal profile in RPKM for TRIs, TRITSSs, and cTSSs. *All profile plot analyses are performed in mBCs. Shaded areas on the unfilled line plots are the standard error.

### Characterization of transcriptional activity at TRIs

The RNAP2s5 signal distribution at TRIs suggests two populations of RNAP2s5, potentially transcribing in opposing directions. To determine if both RNAP2s5 peaks correlate with RNA production, we analyzed nascent transcription at TRIs and cTSSs using a published EU-Seq dataset (69). We found overlapping plus and minus strand nascent transcription signal at TRIs and cTSSs, with more transcription in both directions along TRIs than cTSSs (Figure 2b). These results indicate TRI genes have more bidirectional transcription producing overlapping and partially antisense RNAs. When evaluating full genes, nascent transcription was higher at TRI genes near the TSS (Supplementary Figure S2b, upper panel). Analysis of nascent transcription using an independent GRO-Seq dataset similarly showed overlapping bidirectional transcription at TRIs (73) (Supplementary Figure S2c, upper panel). TRITSSs exhibited more nascent transcription than cTSSs by EU-Seq, while TRITSS and cTSS had similar levels by GRO-Seq (Supplementary Figure S2b and c, lower panels). This apparent difference is likely due to the types of transcriptional activity measured; EU-Seq measures active RNAP *in vivo* but requires longer labeling times, while GRO-Seq measures transcriptionally-competent (paused and active) RNAP molecules *in vitro* (90,91). Taken together, these results indicate that TRITSSs have more active RNAP2 molecules than cTSSs.

The bimodal distribution of RNAP2s5 surrounding TRIs suggests transcription initiation from two distinct sites in close proximity. Upon analyzing the arrangement of genes within TRI regions, three distinct patterns emerged: two annotated genes with non-overlapping divergent TSSs, divergently-transcribing gene pairs with overlapping genic regions, and annotated single genes with unannotated divergent transcription (Figure 2c). Using GRO-Seq to call peaks for plus and minus strand transcription within 3 kb of TRITSSs, we found that ~76% of TRI regions contained peaks on both plus and minus strands indicative of bidirectional transcription. Some regions examined also exhibited EU-Seq signal on both plus and minus strands, consistent with GRO-Seq results (Figure 2c).

R-loops are three-stranded nucleotide structures primarily formed co-transcriptionally by nascent RNA hybridizing to template DNA and looping out the non-template DNA strand. DNA:RNA ImmunoPrecipitation (DRIP) using the S9.6 antibody maps R-loops genome-wide in mammalian cells (92). Similar to RNAPs5 signal, both TRIs and TRITSSs had bimodal DRIP-Seq signal, while cTSSs showed a single central peak of lower amplitude (Figure 2d, Supplementary Figure S2d). We observed higher DRIP-Seq signal along the length of the gene body and at TTSs at cTSS genes than TRI-associated genes, consistent with RNAP2s5 signal (Supplementary Figure S2e). Taken together, the bimodal RNAP2s5 signal and high levels of bimodal DRIP signal at TRIs indicates two distinct populations of active RNAP2 producing RNA molecules with a propensity for template association.

### Epigenetic landscape and chromatin modifier association at TRIs

We next examined the chromatin landscape surrounding TRIs and cTSSs using published ChIP-Seq datasets from mouse primary B cells (73). Like RNAP2s5, the chromatin insulator CTCF and cohesin complex member RAD21 had two flanking peaks of signal at TRIs while cTSS exhibit a single central peak signal (Figure 2e, Supplementary Figure S3a). A similar pattern was observed for the histone acetyltransferases (HAT) p300 and GCN5 (Figure 2e, Supplementary Figure S3a). The increase in HAT occupancy was reflected in histone acetylation, where TRIs exhibited higher H4K12Ac, H4K16Ac and H2BK20Ac signal upstream of the transcription start site relative to cTSSs, as well as histone deacetylases HDAC1 and HDAC2 where there was a single peak in the center of cTSSs and two flanking peaks at TRIs (Figure 2e, Supplementary Figure S3b). These results are consistent with two steady populations of active RNAP2 at TRIs.

To further define the epigenetic state of TRI regions, we next analyzed histone methylation patterns (73) also in stimulated WT mouse primary B cells. Areas flanking TRIs exhibited higher levels of H3K79me1/2, H4K20me1, H3K27me2, H3K4me1/2/3 and H3K9me1—all marks associated with active chromatin (Figure 2f, Supplementary Figure S3c). Further, levels of methylation marks associated with silent or repressed genes—H3K9me2/3 and H3K27me3—were lower than controls (Figure 2g, Supplementary Figure S3d). Together, these results indicate that TRI regions exhibit marks of an active chromatin state (93). Of note, TRIs showed higher flanking signals of H3K79me2 and H4K20me1 than cTSSs (Figure 2f, Supplementary Figure S3c). Both marks are associated with non-homologous end joining (NHEJ) -mediated double-strand break (DSB) repair, possibly suggesting that TRIs accumulate DNA damage (94). The enrichment of chromatin modifying enzymes and histone marks correlating with an active transcriptional state upstream and downstream of TRITSSs is consistent with bidirectional transcription.

### Replication Characteristics of TRIs

#### TRIs are enriched for replication initiation and termination zones

To assess the replication characteristics of TRIs, we analyzed OK-Seq which maps Okazaki fragments of replication forks and can determine replication initiation and termination zones as well as their efficiency and directionality (70) (schematic of expected OK-Seq profiles, Figure 3a). Analysis of OK-Seq signals surrounding TRI and cTSS genes (including 10 kb upstream and downstream of TTS) revealed a positive replication fork directionality (RFD) slope upstream of TRIs and cTSS genes indicating the promoter regions of both groups are origin-rich, in contrast to non-transcribed inactive TSSs (iTSSs) (Figure 3b). These results further support the idea that replication preferentially starts near the transcription start sites of genes (69,95). TRI genes have a steeper positive slope and show higher signal than cTSS genes, indicating that TRI promoter regions tend to contain highly localized and more efficient origins. We also found that OK-Seq had a more negative slope through TRI gene bodies, indicating they experience more termination events than cTSS genes. In a 20 kb window of analysis centered at the TSS, there is a modest dip in RFD signal right at TRITSSs which may indicate these genes contain difficult to replicate areas (Supplementary Figure S4a). These results indicate regions upstream of TRIs are enriched for origins with strong firing efficiency.

**Figure 3.**
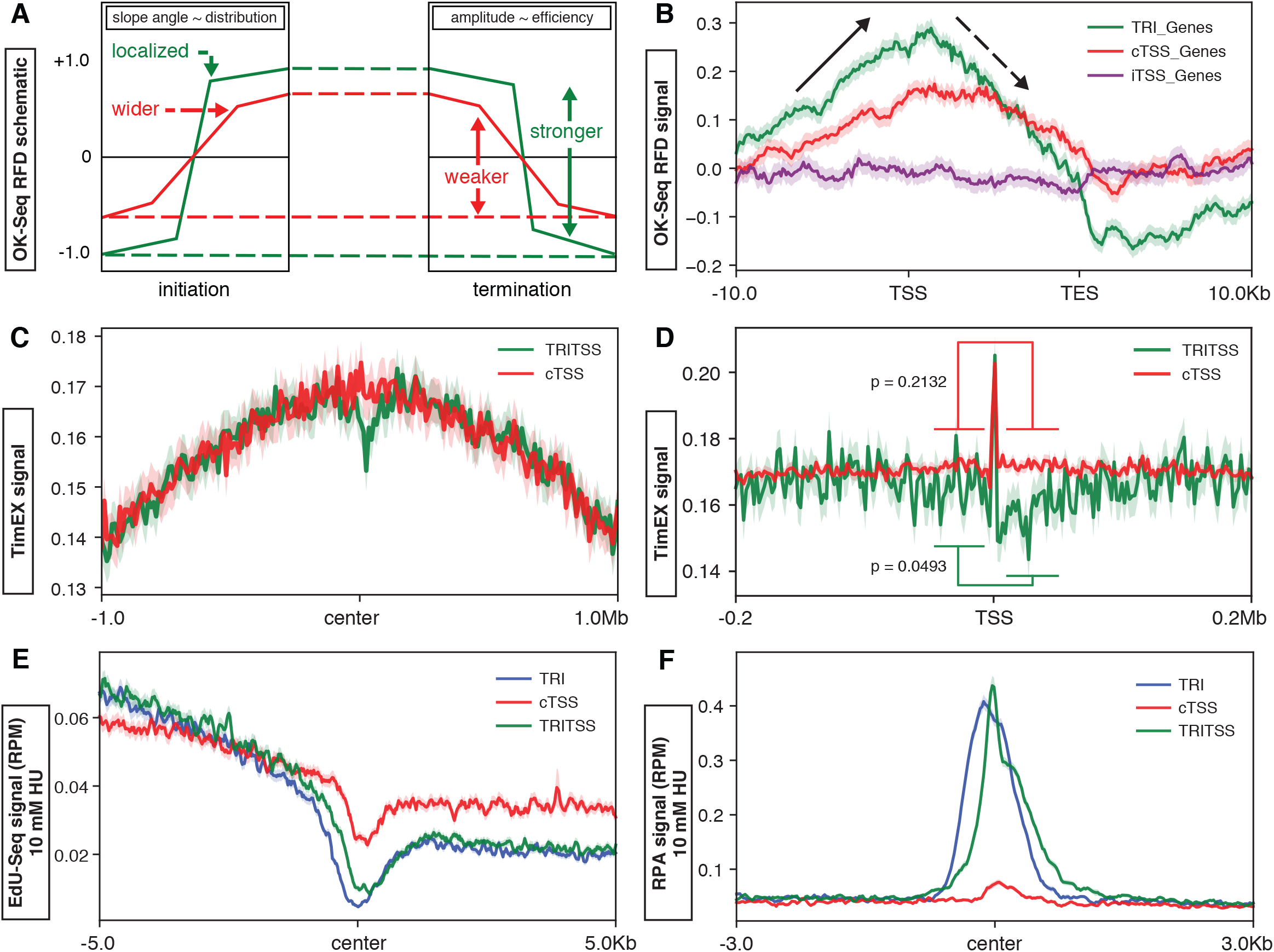
Replication characteristics at TRIs. **(A)** Graphic representation of example of OK-Seq RFD data and interpretation at replication initiation and termination zones: slope correlates with origin localizartion, and amplitude correlates with origin efficiency. **(B)** Median OK-Seq signal profile scaled to full genes +/− 10 kb of TRI genes (green, n=583), cTSS genes (red, n=6,558), and inactive TSS genes (purple, n=11,612). Solid black arrows indicate initiation zones and dashed black arrows indicate termination zones. **(C)** Median TimEX signal profiles centered at TSS +/− 1.0 Mb for TRI genes (green) and cTSSs genes (red). **(D)** Median TimEX signal profiles centered at TSS +/− 200 Kb for TRI genes (green) and cTSSs genes (red). **(E)** Median EdU-Seq signal profiles on cells treated with 10 mM HU centered at TSSs for TRIs, TRITSSs, and cTSSs. **(F)** Median RPA ChIP-Seq signal profiles on cells treated with 10 mM HU centered at TSSs for TRIs, TRITSSs, and cTSSs For TRIs or TRITSS compared to cTSS, p < 1.0 x 10^−250^; Wilcoxon rank sum test). *All profile plot analyses are performed in mBCs. Shaded areas on the unfilled line plots are the standard error.

#### TRIs occur in early replicating zones

To independently assess the replication timing of TRIs, we next analyzed Timing Express (TimEX) data from mouse primary B cells (69). TimEX measures replication timing by calculating the ratio of DNA copy number of cycling cells to resting cells in G0/G1; a higher ratio indicates more replicated DNA in S phase cells and thus earlier replication (96). Analysis of TimEX signal in a 2 Mb window around TRI and cTSS genes shows high TimEX signal, indicating these are some of the earliest replicating regions in the genome (Figure 3c) (69). This is consistent with prior reports showing high transcription correlates with early replication (97). However we observed a reduction in TRITSS TimEX signal near the center, suggesting this region replicates later. Indeed, in a 0.4 Mb window we found that TimEX signal was reduced at TRITSSs, and this reduction was only observed downstream of the TSSs (Figure 3d; p = 0.0493). Unlike TRIs, the TimEX signal for cTSSs was similar upstream and downstream of the TSS (p = 0.2132; Figure 3d). These results indicate that the region downstream of TRITSSs exhibit delayed replication.

#### Contribution of transcriptional activity to replication timing at TRIs

High transcriptional activity has been associated with high origin density, origin efficiency, and earlier replication timing (70). To determine if high transcriptional activity of TRIs can explain their origin enrichment and replication timing, we compared TRIs to cTSSs matched for activity using EU-Seq data. We found that the OK-Seq RFD of TRIs still had increased amplitude and slope compared to transcription activity-matched cTSSs, similar to total cTSSs (Supplementary Figure 4b). Thus, transcriptional activity alone does not explain the observed origin enrichment and early replication timing observed at TRITSSs.

#### TRIs are located near early replication origins

OK-Seq and TimEX define both TRIs and cTSSs as origin-rich and early replicating. To confirm these results, we assessed DNA replication using EdU-Seq analysis (69). EdU-Seq assesses early S phase origin firing by stimulating G0 splenic B cells to enter S phase in the presence of hydroxyurea (HU) to slow or stop replication forks and the thymidine analog (5-ethynyl-2’-deoxyuridine) EdU to label nascent DNA (98); hence, a high EdU signal implies early replication. We found that EdU-Seq signal in HU-treated cells is highest upstream of the TSSs with strong depletion of signal at the center and an increase in signal after the TSSs (Figure 3e). The pattern at TRIs and TRITSSs was more pronounced than cTSSs. These results suggest that TRIs initiate replication earlier than cTSSs but take longer to complete replication downstream; this interpretation is supported by TimEX analysis (Figure 3c,d). We observed similar patterns using EdC-Seq—a variant of EdU-Seq using a cytosine analogue for labeling— from HU-treated cells, indicating that this pattern is independent of sequence (Supplementary Figure S4c) (69). We also observed a dip EdU-Seq signal at TRIs and cTSSs in the absence of HU (Supplementary Figure S4d). From this data, we propose that TRIs experience frequent replication fork stalling with our without exogenous stress.

#### TRIs exhibit high levels of Replication protein A in cells experiencing replication stress

Stalled replication forks accumulate single-stranded DNA (ssDNA) that can be bound by replication protein A (RPA) which then acts as a signal for other processes to repair the stalled replication forks (99). In HU-treated cells, TRIs and TRITSSs exhibit significantly more RPA signal than cTSSs with the signal centered at the TSSs (p = < 1.0 x 10^−250^, Figure 3f). In yeast, RPA accumulates on the lagging strand during HU-induced replication stress (100). To determine if RPA association shows a similar strand bias in mammalian cells, we next measured RPA signal along plus and minus strand surrounding TRIs. For both populations, we observed RPA signal was higher on the lagging strand upstream of the TRI when treated with HU, consistent with studies of stalled replication forks in yeast (Supplementary Figure S4e) (101).

#### TRIs are enriched for Early replicating fragile sites

It is hypothesized that chromosomal fragile sites—genomic regions experiencing recurrent DNA breaks due to replication stress—may be a result of transcription-replication collisions (102). We analyzed the association of TRIs with 615 early-replicating fragile sites (ERFSs) encompassing 126,624 kb (61) and 17 late-replicating common fragile sites (CFSs) encompassing 167,578 kb (62–67). Fragile sites are very large genomic regions ranging from 50 kb to over 2 Mb while the median TRI length was 1,180 bp, so we investigated the overlap in basepairs instead of the number of fragile sites and TRIs. There was a 2.8-fold enrichment of TRIs in ERFS-associated regions, but no enrichment of TRIs in CFS regions (Supplementary Figure S4f, g). ERFSs are enriched for gene pairs exhibiting divergent transcription visible as two TSSs within 3 kb of each other (61). Similar to ERFSs, we found TRITSSs had divergently-transcribing gene pairs, such as *Thap4/Atg4b* (Figure 2c). The enrichment of TRIs for ERFSs but not CFSs is consistent with OK-Seq and TimEX analyses indicating TRIs are origin-rich and early replicating.

### DNA double-strand break formation at TRIs

#### Spontaneous DSBs are enriched at TRIs

TRIs are putative areas of transcription-replication conflicts and potential sites of stalled replication forks and DNA DSBs (103). To assess if TRIs are enriched for DSBs, we analyzed published END-Seq datasets which map exposed DNA ends genome-wide in murine B cells (58). Over 87% of TRIs overlapped with END-Seq signal peaks, a 33-fold enrichment over random chance (Supplementary Figure S5a). To define where DSBs occur relative to transcription and replication at TRIs, we compared END-Seq NT, EU-Seq NT, and EdU-Seq NT datasets from non-treated cells harvested 28 h post-stimulation. Signal intensities were adjusted to fit all three sample types on the same scale. We found that the EdU-Seq signal decreased at it approaches TRIs, while EU-Seq signal sharply increased at the center of TRIs (Figure 4a). END-seq signal was highest slightly upstream of where replication and transcription coincide at TRIs. From this data we conclude that TRIs are enriched for spontaneous DSBs immediately upstream of where transcription begins and replication timing delays occur.

**Figure 4.**
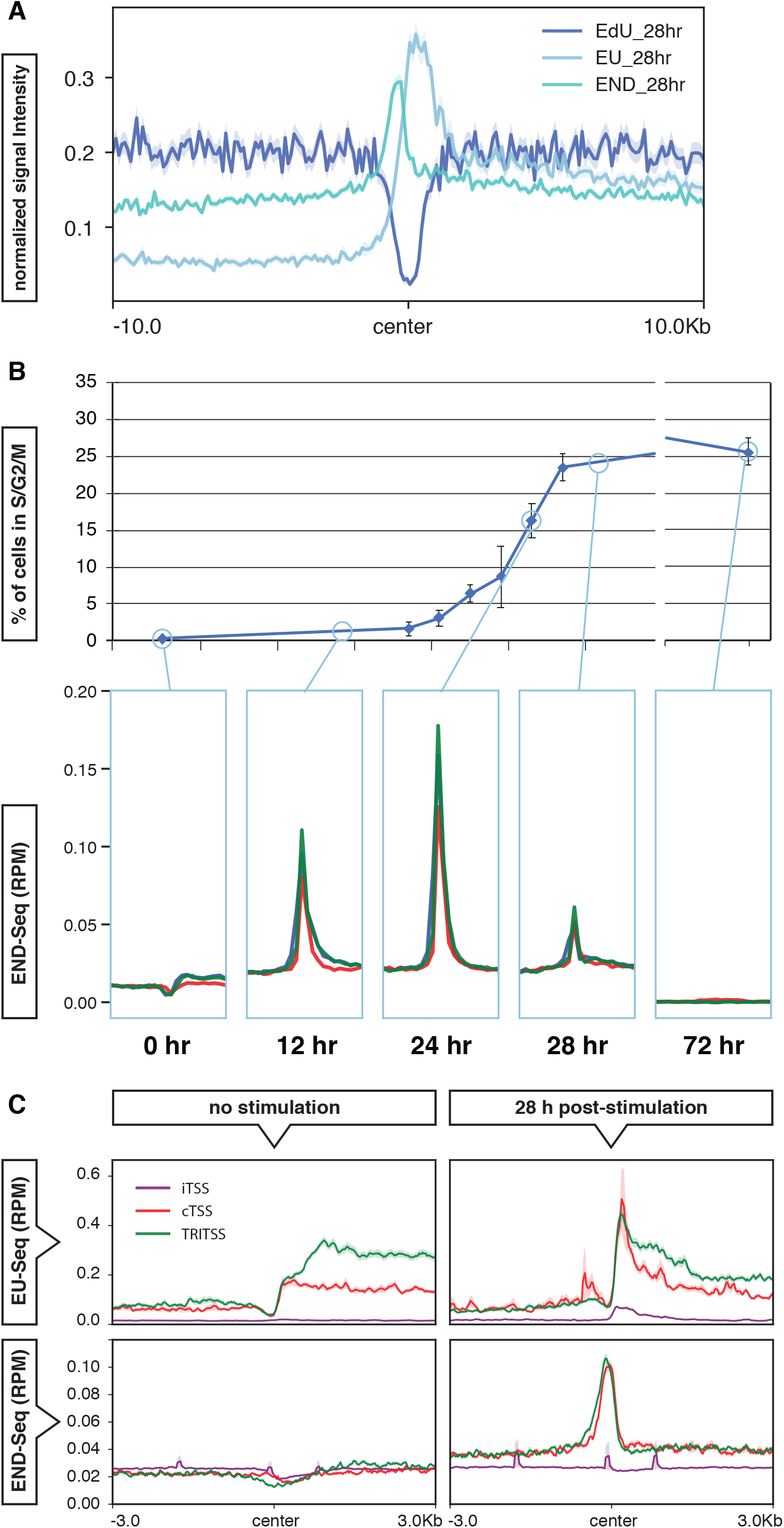
DNA breaks at TRIs. **(A)** Median signal profiles centered at TRIs of EdU-Seq (blue), EU-Seq (dark blue), and END-Seq (cyan) from 28 hour NT cells. The EdU-Seq and END-Seq signal was multipled by 8 and 4 respectively so the signals can be seen clearly on the same scale. **(B)** Top graph showing the percentage of B cells in S and G2/M phases measured by PI incorporation and analyzed by FACS. Bottom of the panel showing END-Seq signal profile at 0, 12, 24, 28, and 72 hour time points from NT cells centered near TSSs of TRIs (blue), TRITSSs (green), and cTSSs (red). **(C)** Median signal profiles of EU-Seq (top) and END-Seq 0 mM HU (bottom) of non-replicating resting (left) and 28 hour activated replicating mouse B cells (right) centered near TSSs of TRITSSs (green), cTSSs (red), and inactive TSSs (purple). *All profile plot analyses are performed in mBCs. Shaded areas on the unfilled line plots are the standard error.

#### Spontaneous DSBs correlate with transcription and replication activity at TRIs and cTSSs

We next compared the END-Seq signal at TRIs and cTSSs with entry into S phase to determine if DSB accumulation correlates with DNA replication. In response to antigen stimulation, naïve splenic B cells resting in G0 first undergo an increase in transcription, then enter the cell cycle and replicate (104). Flow cytometric analysis of DNA content shows that DNA replication initiates around 14-16 hours post stimulation, with ~16% of cells in S phase by 24 hours (Figure 4b, top). END-Seq signal positively correlated with cells entering S phase for both TRIs and cTSSs, indicating these DSBs are replication-dependent and do not simply represent transcription-induced damage (Figure 4b, bottom) (58,69,105).

To determine if both replication and transcription are required for DSB formation, we compared END-Seq and EU-Seq data from replicating and non-replicating cells at TRITSSs, cTSSs and inactive TSSs (iTSSs) (58,69). We found EU-Seq signal at TRIs and cTSSs in non-replicating cells (Figure 4c, upper left) but no enrichment of END-Seq signal (Figure 4c, lower left). In replicating cells, we found END-Seq signal accumulation at actively-transcribing TRIs and cTSSs, but not iTSSs which had minimal transcription by EU-Seq (Figure 4c, right). These results suggest that DSBs formed at TRIs and cTSSs accumulate only when replication and transcription are both active. The Ataxia telangiectasia and Rad3-related checkpoint kinase (ATR) plays a critical role in the repair and/or restart of stalled replication forks (61,106). To determine if loss of ATR enhanced DNA damage at TRIs, we analyzed END-Seq data at TRIs and cTSSs in the absence and presence of the ATR inhibitor (ATRi) AZ20. We observed an increase in the END-Seq signal in cells treated with 10 μM ATRi at both TRIs and cTSSs, consistent with the notion that ATR helps suppress DNA damage arising at these loci (Supplementary Figure S5b) (69).

#### DSBs form independently of TOP2

Topoisomerase II (TOP2) relieves topological stress through its DNA cleavage-religation activity; inhibition of TOP2 re-ligation with etoposide leads to single-strand breaks (SSBs) and DSBs (107). To assess if TOP2 activity contributes to DSB formation at TRIs, we analyzed END-Seq datasets for etoposide-treated WT and TOP2B knockout (TOP2BKO) mouse B cells as a function of time post B cell simulation (58). When non-replicating WT cells were exposed to etoposide, END-Seq signal increased in regions flanking the TSS but not at the start site itself, suggesting these DSBs are associated with transcription only (Supplementary figure 4c). When replicating WT cells were treated with etoposide, the END-Seq signal again accumulated in the regions flanking TRI/TRITSSs with no noticeable increase in the center; however, in TOP2BKO cells the END-Seq signal was still centered at the TSS (Supplementary Figure S5c) (108). These results suggest that the flanking DSBs forming around TRIs are associated with transcription and TOP2B activity, while DSBs directly at the center of TSSs are largely TOP2B-independent.

To investigate how DSBs are distributed around TRIs, we analyzed 24 hr END-Seq signal on the positive and negative DNA strands. We found that the END-Seq signal at TRIs was elevated and had a wider distribution than cTSSs, suggesting that DNA breakpoints are more localized at cTSSs (Supplementary Figure S5d). In previous studies, END-Seq signal accumulated unevenly around poly(dA:dT) tracts in response to HU with a ratio of ~2:1, suggestive of fork collapse from a stalled DNA polymerase (69). However, the distribution of END-Seq signal at TRIs was 1:1; therefore, we speculate that polymerase stalling is not the dominant cause of TRI-associated DSBs.

### Sequences and mutations at TRIs

#### GC sequences are enriched at TRIs

Repetitive DNA elements such as trinucleotide repeats and long inverted repeats can form secondary structures capable of inducing replication fork blockage (109). We examined the nucleotide content and found that TRITSSs are significantly more GC-rich than cTSSs, with enrichment peaking at TRI centers (p < 1.0 x 10^−250^, Figure 5a; Supplementary Figure S6a). Additionally, 99% of TRIs overlap with CpG islands, a 78-fold enrichment over random chance (Supplementary Figure S6b). CpG islands correlate with low DNA methylation levels (110). Consistent with this notion, TRITSSs had significantly lower DNA methylation than cTSSs (111) (p = 8.3 x 10^−238^, Figure 5b). We next measured GC-skew, an asymmetrical distribution of nucleotides where guanines are more abundant than cytosines. GC-skew has been associated with R-loops, CpG island promoters, and prokaryotic replication origins (78,112). We found a sharp increase to a positive GC-skew at the center of both TRITSSs and cTSSs; however, TRITSSs exhibited significantly higher GC-skew than cTSSs within gene bodies (p = 1.48 x 10^−67^, Figure 5c). G-quadruplex (G4) motifs can form secondary structures when guanine-rich sequences form a helical shape stabilized by G-G base pairing and have been implicated in genome instability (113). We used G4 Hunter to predict G4 structure formation, and found that 93% of TRIs and 70% of cTSSs can potentially form G4s (77). TRIs average 1 possible G4 per 104 bp, while cTSSs could form 1 possible G4 per 149 bp (Figure 5d). Using Homer, we also found an enrichment of two similar GC-rich sequences, CCGCCGCC and GGCGGCGG, in both TRIs and cTSSs but the frequency was higher in TRIs. (Supplementary Table S3). These results show that TRIs are enriched for GC content, potential secondary structure-forming G4 sequences, and GGC trinucleotide repeats.

**Figure 5.**
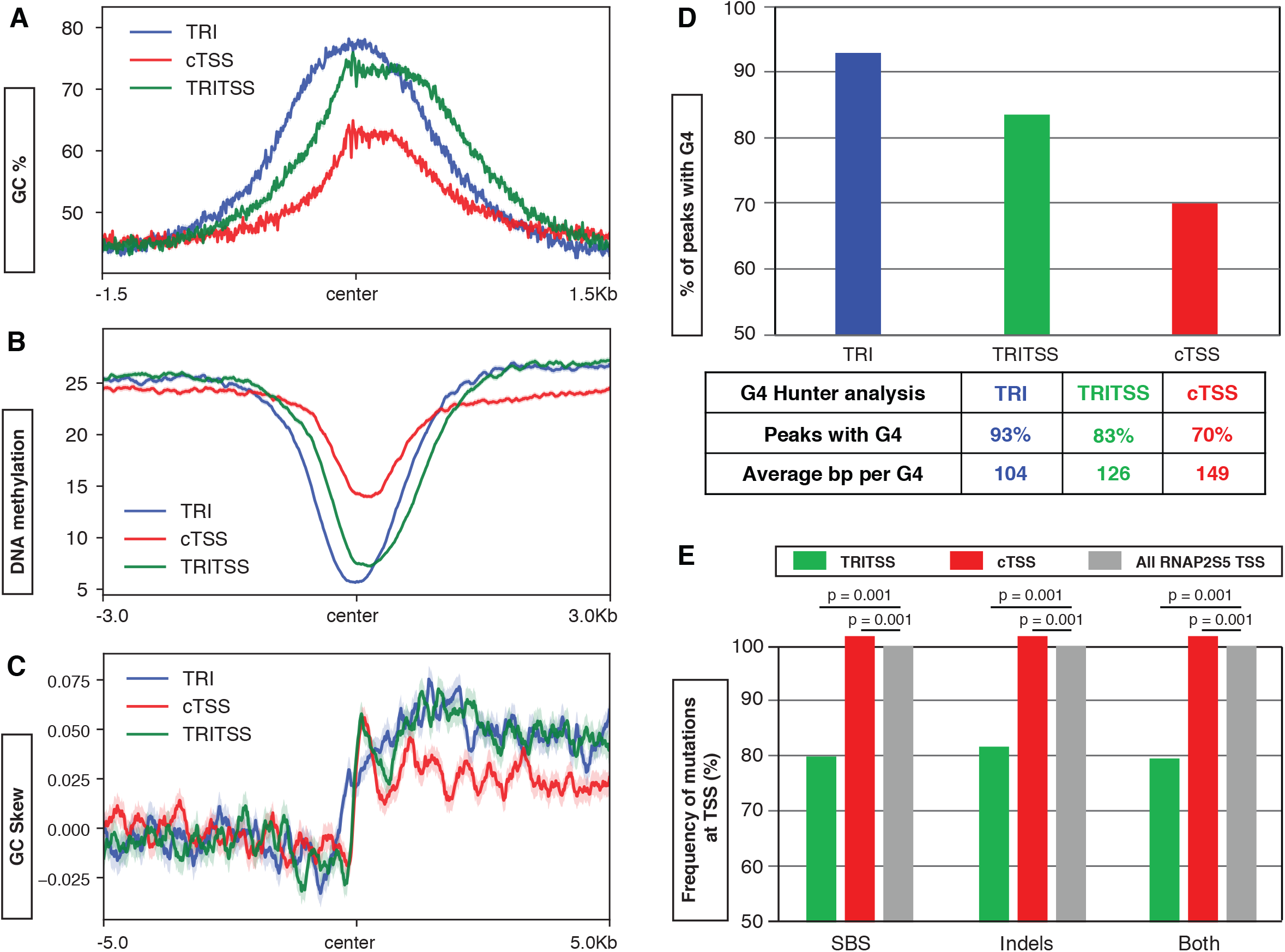
Sequences and mutations at TRIs. **(A)** Median signal profile of percent GC of the mm10 mouse genome centered near the TSSs of TRIs (blue), TRITSSs (green), and cTSSs (red). **(B)** Median DNA methylation signal profile of mouse B cells centered near the TSSs of TRIs (blue), TRITSSs (green), and cTSSs (red). For TRIs vs. cTSS, p < 1.0 x 10^−250^, and TRITSS vs. cTSS, p = 8.3 x 10^−238^; Wilcoxon rank sum test. **(C)** Median signal profile of GC skew of the mm10 mouse genome centered near the TSSs of TRIs (blue), TRITSSs (green), and cTSSs (red). GC skew of TRIs and TRITSS are significantly higher than cTSS (p = 1.17 x 10^−66^ and p = 1.48 x 10^−67^ respectively, Wilcoxon rank sum test). **(D)** Table and bar graph of the predicted frequency of G-quadruplex formation at TRIs, TRITSSs, and cTSSs. **(E)** Bar graph of single basepair substitutions (SBS) and short insertions/deletions (indels) at TRIs, TRITSSs, and cTSSs normalized to cTSSs showing empirical p-value. *Shaded areas on all unfilled line plots are the standard error.

#### TRIs accumulate deletion mutations

TRIs and cTSS are enriched for DNA breaks by END-Seq; therefore, it is possible they accumulate mutations. We analyzed sequence variants containing single basepair substitutions (SBSs) and short insertions/deletions (indels) in four independent datasets from mouse primary B cells—two TimEX sequencing results and two datasets used as input controls for ChIP-Seq experiments (69,73). Compared to all RNAP2S5-associated TSSs, TRITSSs harbored fewer SBSs and indels (p < 0.001, Figure 5e). However both mutation types were increased at cTSSs (p < 0.001, Figure 5e). Upon further analysis of indels, we found an enrichment of deletions (Supplementary Table S4). Together, these results indicate that TRITSSs accumulate fewer mutations than cTSSs, but TRI gene bodies accumulate more mutations than cTSS genes.

The Catalogue Of Somatic Mutations In Cancer (COSMIC) has developed a set of bioinformatic tools to perform variant analysis and identify specific mutational signatures from next-generation sequencing data (114). To investigate the specific types of mutations occurring at TRIs and cTSSs, we analyzed the SBSs and indels using the COSMIC SigProfiler software suite. We found an overrepresentation of C(C>A)G mutations in TRITSS regions and full genes compared to cTSSs (Figure S6d - blue bars, Supplementary Table S4). TRITSSs also had more 3 bp deletions at repeats than cTSSs (Supplementary Figure S6e - pink/orange lines, Supplementary Table S4). These distinct mutations may occur because TRIs have more trinucleotide CCG repeats therefore they are overrepresented, or because CCG/GGC motifs can cause DNA polymerase stalling (115). Thus, though mutations are less frequent in TRITSS, they preferentially accumulate small deletions in repetitive DNA sequences.

#### TRI genes overlap with cancer drivers

We next probed a dataset from the Mouse Tumor Biology Database (MTBD) that listed 23,625 mutations (6,525 unique gene names) from sequenced mouse tumors (71). We found that 57.3 % of TRI genes harbor mutations, a 3.4 fold enrichment over random chance (Supplementary Figure S6f). Insertions were overwhelmingly the most abundant mutation in the MTBD dataset at 73% of all mutations (Supplementary Table S5). When analyzing MTBD mutations at TRIs we observed an underrepresentation of insertions (79%) and an overrepresentation of deletions (227%), nonsense mutations (356%), and point mutations (214%) (Supplementary Table S5, Supplementary Figure S6g). Since TRI gene mutations associate with mouse tumors, we next explored the Sleeping Beauty Cancer Driver DataBase (SBCDDB) which identified 1,231 cancer drivers (72). Here we observed an 29% overlap with SBCDDB cancer drivers, a 9.2-fold enrichment over random genes. These results show TRI regions accumulate specific mutation subtypes in murine tumors.

#### TRI gene set enrichment analysis

DNA damage and accumulation of somatic mutations have been hypothesized to be involved with cancer (116). Using MouseMine, we found that TRI genes are highly enriched for preweaning lethality, abnormal survival, mortality, and aging phenotypes in mice (Supplementary Table S6) (80). We found that TRI genes were enriched in KEGG pathways associated with cancer and transcription factor protein-protein interactions (PPIs) with cancer associated genes such as *Tp53*, *Brca1*, and *Myc* (Supplementary Table S7). Our results also showed an association of TRI genes with stem cell pluripotency factors; pluripotency factors can be induced in cancers and associate with poor treatment outcomes (Supplementary Table S8) (117). This association of TRIs with cancer is supported by the overlap between MTB tumor mutations and SBCDDB cancer drivers. Chromatin structure, post-translational modifications, and gene expression have been implicated in aging and cancer. We also observed an enrichment of TRI genes in these processes as well as PPIs with *Ep300* and *Hdac2* (Supplementary Table S9 (118).

## DISCUSSION

In this study we developed a method to identify genomic locations where transcription and replication machinery colocalize genome-wide, identifying 1,009 independent TRIs in primary mouse B cells. TRIPn-Seq can be applied to any proliferating cell type, as it relies on incorporation of modified nucleotides into nascent replication. A subset of TRIs overlap two unique annotated genes, therefore we identified TRIs at 1,198 active genes. However, these results do not rule out transcription-replication problems at the other 12,957 active genes. Rather, TRIs may represent areas of prolonged or complex interactions. Indeed, the bidirectional transcription so prevalent at TRIs strongly increases the chance of conflicts with replication machinery in proliferating cells.

TRIs harbor distinct patterns of chromatin features which are distinct from other transcribed genes. In particular, TRIs are characterized by a bimodal pattern for RNAP2s5, R-loops, P300, GCN5, HDAC1 and HDAC2. Transcriptional activity influences the placement of proteins involved in chromatin architecture; indeed, we see a similar bimodal pattern in RAD21 and CTCF at TRIs. From the RNAP2s5 ChIP-Seq libraries generated here we cannot confirm the simultaneous binding or direction of two RNAP2s5 molecules on the same DNA strand, but the presence of two RNAP2s5 populations is supported by the bidirectional overlapping transcription measured by EU-Seq and GRO-Seq (Figure 2b, c and Supplementary Figure S2b,c). This transcriptional landscape indicates that multiple RNA polymerase complexes are engaged on the template and non-template strand surrounding TRITSSs. Multiple RNAPs moving in a codirectional or convergent orientation with respect to replication are more difficult impediments to replication than a single RNAP, and may explain why TRIs were detected at these locations (16). Divergent transcription has also been shown to increase genome instability and interactions with replication machinery likely exacerbates this (119). Although transcription and replication machineries may interact at all transcribed loci, it is possible that only a specific orientation of RNAP2s5 causes prolonged or increased frequency of fork stalling at TRIs. Overall, TRI genes show higher levels of nascent transcription relative to cTSS genes, as well as chromatin marks associated with active transcription (Figure 2b, Supplementary Figure S3). Thus, higher levels of RNAP sense transcription may increase interactions with ongoing replication, increasing TRI frequency.

Intriguingly, TRIs and TRITSSs also exhibited a bimodal pattern for DNA:RNA hybrid formation more similar to RNAP2s5 ChIP-Seq signal than active transcription as measured by EU-Seq or GRO-Seq where the upstream signal is low. GC skew has been associated with R-loop formation, yet we only observed GC skew downstream of TRITSSs (112). This begs the question, what stimulates R loop formation upstream of TRITSSs? One possibility is that upstream R loops are stimulated by RNAP2s5 binding. Paused RNAP2 could anchor nascent RNAs in place, promoting interaction with the template strand. Recent studies also suggest that DNA supercoiling can drive RNA:DNA hybrid formation in sequences without significant GC skew; this may relieve torsional stress by allowing the DNA strand to twist around the RNA (120). Thus, regions upstream of TRITSSs forming R-loops may “absorb” negative supercoiling, while GC skew promotes R-loop formation downstream of the TRITSS. Both transcription and replication generate negative supercoiling behind elongating complexes and positive supercoiling in front suggesting a reliance on topoisomerases to maintain genomic integrity. Intriguingly, analysis of END-Seq data from etoposide-treated TOP2BKO cells show that DSBs still accumulate at the TSSs, even though flanking DSBs decrease from WT (Supplementary Figure S5c) (107,121). This evidence may imply that TOP2B is not the sole enzyme creating DSBs at TRIs; in the absence of TOP2B, additional enzymes may process topological stress at these sites such as TOP1 as shown in recent studies (122).

Genetic and epigenetic signatures of TRIs point to a chromatin landscape providing multiple roadblocks to efficient replication, leading to fork stalling and collapse. This is reflected in OK-Seq analyses which show a distinct pattern of replication around TRIs with origins enriched upstream of the TSSs, and a strong increase in termination throughout gene bodies and TTSs (Figure 3a). This pattern is consistent with prior publications showing replication termination is enriched within gene bodies in response to HU-induced replication stress (95). Regions upstream of TRIs appear to be some of the earliest replicating loci in the genome as measured by EdU-Seq, EdC-Seq and TimEX experiments. Early replicating regions are associated with high transcription, increasing the chance of transcription-replication conflicts in early S phase (123). All replication datasets examined show subtle but consistent dips in replication efficiency at the center of TRIs(Figure 3b-e, Supplementary Figure S4a-d). Similar studies measuring DNA replication and transcription throughout S phase in human fibroblasts also show a strong delay in replication around TSSs (124). Thus, we hypothesize TRIs are difficult-to-replicate regions enriched for RF stalling and collapse. Indeed, TRIs accumulate extensive RPA—a hallmark of replication fork stalling and DSB formation (125).

We propose that TRIs represent genomic loci which can stall both the replicative helicase and DNA polymerase. In our model, replication initiates upstream of active genes. Near the TSS of TRIs, the helicase of a moving replisome may stall on the leading strand, while the lagging strand DNA polymerase is can stall at a complex GC sequence or DNA:RNA hybrid (Figure 6a). A double strand break is formed and the RNAP2s5, replication fork, or both are displaced. There is a short amount of resection (< 500 bp), RPA is loaded onto the ssDNA, and the damage is repaired by microhomology-mediated end joining (MMEJ). How is replication at TRIs completed if they are such roadblocks to fork progression? Replication across TRIs can be completed by fork restart if the replisome remains associated, or by passive replication from adjacent origins (Figure 6b). Another possibility is that TRIs are bypassed and replicated later through a non-conventional replication mechanism. In agreement with our findings, recent evidence indicates that a subset of TSSs are replicated in G2/M (124). EdU-Seq also provides some evidence for this possibility as downstream signal is higher than at the center (Figure 3c). Alternatively, downstream replication forks may replicate this area resulting in fork termination at TRIs (Figure 6b).

**Figure 6.**
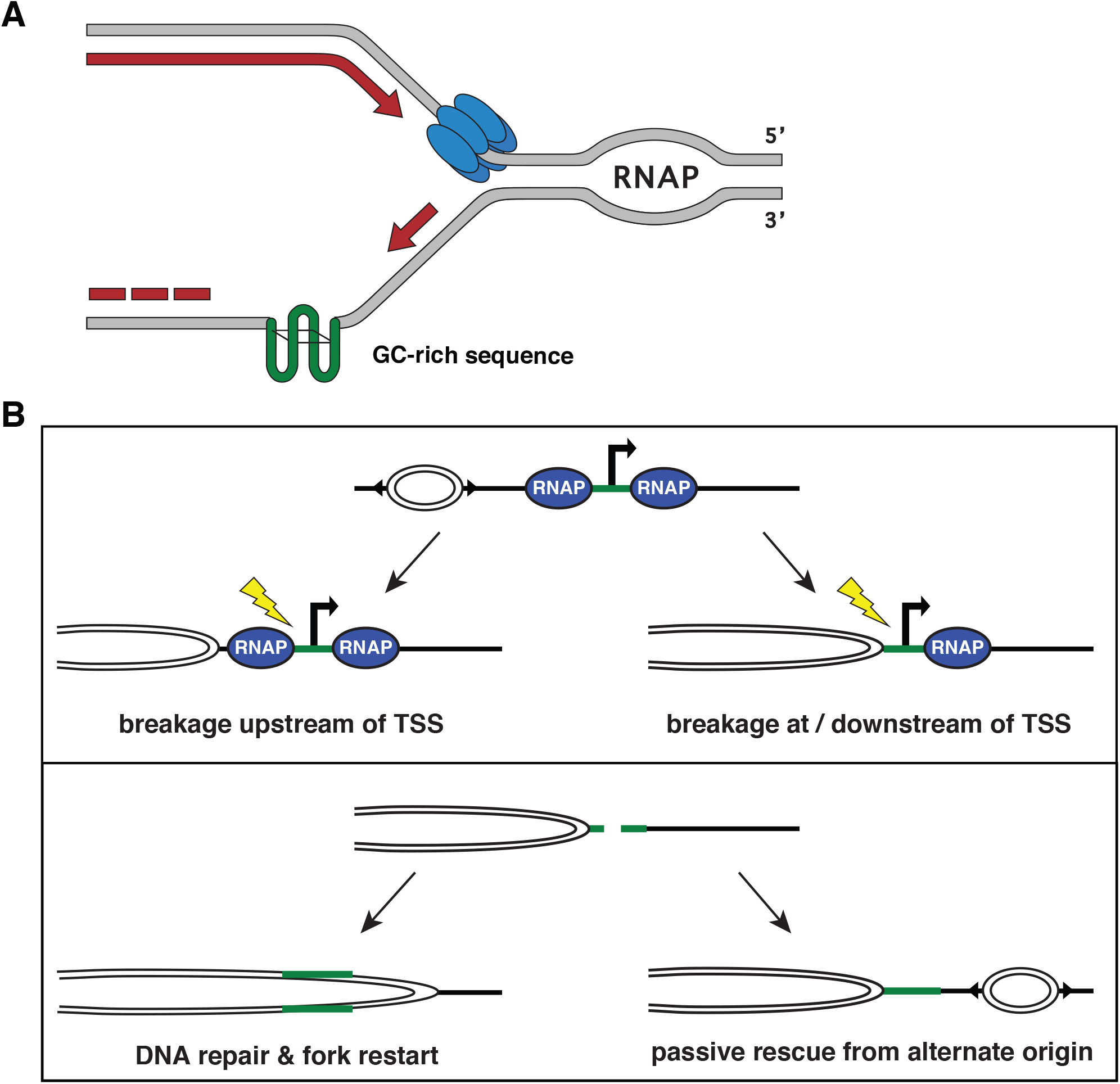
Summary of features at TRIs and model of TRIs. **(A)** Graphic summary of chromatin features at TRIs and chromatin features of cTSSs. **(B)** Graphic representation of a TRI where the replicative helicase is stalled by an RNAP molecule and the DNA polymerase is stalled by a complex GC-rich sequence or structure (green) simultaneously.

DSBs need to be repaired to preserve genome integrity and TRIs show higher flanking signals of H4K16Ac, H3K36me2, H3K79me2, and H4K20me1 all of which are involved in non-homologous end joining (NHEJ) during DSB repair when compared to cTSSs (Supplementary Figure S3) (94). TRIs also have considerably more RPA in HU treatment than cTSSs indicating persistent ssDNA formation. This accumulation of RPA-bound ssDNA may be a hallmark of DSB repair by homologous recombination (HR) at TRIs or simply an accumulation of stalled replication forks (99). Alternatively, RPA is also present at sites of alternative end joining involving ssDNA tails which contributes to deletions and translocations (126).

TRIs are enriched for small deletions, consistent with the increase in END-Seq signal and NHEJ-associated chromatin marks. TRIs have a wider distribution of DSBs compared to cTSS which may imply that TRI-associated DSBs are generated by alternate means, or the DSB ends are processed more extensively for repair. Processing of DSB ends revealing ssDNA would make TRI breaks substrates for MMEJ, a mutagenic form of NHEJ with a propensity for inducing microdeletions (127). Indeed, MMEJ repairs breaks at collapsed replication forks, and convergent transcription-replication interactions increase the frequency of deletion events (128,129). The enrichment of trinucleotide repeats at TRIs are also likely to play a role in the increase in deletions, as MMEJ requires short regions of homology. Replication slippage also occurs at trinucleotide repeats, resulting in small deletions as well as expansions of repeated sequences and requires disruption of continuous replication. Our mutational signature analysis shows that there are patterns that match COSMIC samples experiencing replication slippage (Supplementary Figure S6e) (115,130). It is therefore surprising that we find fewer single nucleotide variants at TRIs than cTSS, since they harbor characteristics of MMEJ (131). Though TRIs have elevated DSBs by END-Seq and high levels of RPA, TRI genes are depleted for single basepair mutations and indels. One possibility is that TRIs experience less mutagenic DNA repair. TRI genes are highly enriched for preweaning lethality, abnormal survival, mortality, and aging phenotypes. Thus mutations in TRI genes may induce cell lethality, leading to fewer mutations in sequencing analysis. Further experiments assessing DSB repair pathway choice at TRIs will define the contribution of NHEJ, HR and alternative repair pathways at these sites.

TRIs may also have a role in promoting genome integrity. Codirectional transcription-replication collisions can induce replication fork restart and activate the DNA damage response (27,132). Thus, TRI-induced fork pausing at TSSs may minimize co-directional collisions from occurring within the coding regions of essential genes. This is similar to replication fork barriers that are found in active ribosomal DNA termination regions which prevent convergent TRIs within the gene (133). If the cell survives, acquired mutations can persist and potentially accumulate leading to aging phenotypes. Changes near TSSs are more likely to induce expression changes which could eventually reach a critical limit resulting in gene silencing or overexpression. At TRI genes such as *Cop1*, a negative regulator of tumor suppressor gene *Tp53*, this could disrupt the balance and drive tumor formation. Mechanistic studies investigating TRI-associated DSB formation and repair will help untangle how transcription and replication influence genome instability, tumorigenesis, and aging.

## DATA AVAILABILITY

TRIPn-Seq, DRIP-Seq, and ChIP-Seq data has been deposited to the Gene Expression Omnibus (GEO) database under the accession GSE161410 and the Flow Repository under ID# FR-FCM-Z3GS. Previously published data analyzed in these experiments are from GSE100262 (111), GSE116321 (69), GSE129524 (108), GSE99197 (58), GSE82144 (73), and SRA: PRJNA326246 (105). mm10 GC percent (60)

## ACKNOWLEDGEMENTS

We would like to thank Drs. Wolf-Dietrich Heyer and Duncan Smith for critical reading of the manuscript.

## REFERENCES

1. Macheret, M. and Halazonetis, T.D. (2015) DNA replication stress as a hallmark of cancer. Annu Rev Pathol, 10, 425–448.

2. Kim, N. and Jinks-Robertson, S. (2012) Transcription as a source of genome instability. Nat Rev Genet, 13, 204–214.

3. Helmrich, A., Ballarino, M. and Tora, L. (2011) Collisions between replication and transcription complexes cause common fragile site instability at the longest human genes. Molecular cell, 44, 966–977.

4. Helmrich, A., Ballarino, M., Nudler, E. and Tora, L. (2013) Transcription-replication encounters, consequences and genomic instability. Nat Struct Mol Biol, 20, 412–418.

5. Gomez-Gonzalez, B. and Aguilera, A. (2019) Transcription-mediated replication hindrance: a major driver of genome instability. Genes Dev, 33, 1008–1026.

6. Hsin, J.P., Xiang, K. and Manley, J.L. (2014) Function and control of RNA polymerase II C-terminal domain phosphorylation in vertebrate transcription and RNA processing. Molecular and cellular biology, 34, 2488–2498.

7. Nojima, T., Rebelo, K., Gomes, T., Grosso, A.R., Proudfoot, N.J. and Carmo-Fonseca, M. (2018) RNA Polymerase II Phosphorylated on CTD Serine 5 Interacts with the Spliceosome during Co-transcriptional Splicing. Molecular cell, 72, 369–379 e364.

8. Guo, J. and Price, D.H. (2013) RNA polymerase II transcription elongation control. Chemical reviews, 113, 8583–8603.

9. Itoh, T. and Tomizawa, J. (1980) Formation of an RNA primer for initiation of replication of ColE1 DNA by ribonuclease H. Proc Natl Acad Sci U S A, 77, 2450–2454.

10. Gan, W., Guan, Z., Liu, J., Gui, T., Shen, K., Manley, J.L. and Li, X. (2011) R-loop-mediated genomic instability is caused by impairment of replication fork progression. Genes Dev, 25, 2041–2056.

11. Obi, I., Rentoft, M., Singh, V., Jamroskovic, J., Chand, K., Chorell, E., Westerlund, F. and Sabouri, N. (2020) Stabilization of G-quadruplex DNA structures in Schizosaccharomyces pombe causes single-strand DNA lesions and impedes DNA replication. Nucleic Acids Res, 48, 10998–11015.

12. Samadashwily, G.M., Raca, G. and Mirkin, S.M. (1997) Trinucleotide repeats affect DNA replication in vivo. Nat Genet, 17, 298–304.

13. Ait Saada, A., Lambert, S.A.E. and Carr, A.M. (2018) Preserving replication fork integrity and competence via the homologous recombination pathway. DNA Repair (Amst), 71, 135–147.

14. Schroeder, J.W., Sankar, T.S., Wang, J.D. and Simmons, L.A. (2020) The roles of replication-transcription conflict in mutagenesis and evolution of genome organization. PLoS Genet, 16, e1008987.

15. Chong, S.Y., Cutler, S., Lin, J.J., Tsai, C.H., Tsai, H.K., Biggins, S., Tsukiyama, T., Lo, Y.C. and Kao, C.F. (2020) H3K4 methylation at active genes mitigates transcription-replication conflicts during replication stress. Nat Commun, 11, 809.

16. Bruning, J.G. and Marians, K.J. (2020) Replisome bypass of transcription complexes and R-loops. Nucleic Acids Res, 48, 10353–10367.

17. Fritz, A., Sinha, S., Marella, N. and Berezney, R. (2013) Alterations in replication timing of cancer-related genes in malignant human breast cancer cells. J Cell Biochem, 114, 1074–1083.

18. McGlynn, P., Savery, N.J. and Dillingham, M.S. (2012) The conflict between DNA replication and transcription. Mol Microbiol, 85, 12–20.

19. Jonkers, I. and Lis, J.T. (2015) Getting up to speed with transcription elongation by RNA polymerase II. Nat Rev Mol Cell Biol, 16, 167–177.

20. Prado, F. and Aguilera, A. (2005) Impairment of replication fork progression mediates RNA polII transcription-associated recombination. The EMBO journal, 24, 1267–1276.

21. Rocha, E.P. and Danchin, A. (2003) Gene essentiality determines chromosome organisation in bacteria. Nucleic Acids Res, 31, 6570–6577.

22. Pomerantz, R.T. and O’Donnell, M. (2010) Direct restart of a replication fork stalled by a head-on RNA polymerase. Science, 327, 590–592.

23. Paul, S., Million-Weaver, S., Chattopadhyay, S., Sokurenko, E. and Merrikh, H. (2013) Accelerated gene evolution through replication-transcription conflicts. Nature, 495, 512–515.

24. Pomerantz, R.T. and O’Donnell, M. (2008) The replisome uses mRNA as a primer after colliding with RNA polymerase. Nature, 456, 762–766.

25. Sankar, T.S., Wastuwidyaningtyas, B.D., Dong, Y., Lewis, S.A. and Wang, J.D. (2016) The nature of mutations induced by replication-transcription collisions. Nature, 535, 178–181.

26. Dutta, D., Shatalin, K., Epshtein, V., Gottesman, M.E. and Nudler, E. (2011) Linking RNA polymerase backtracking to genome instability in E. coli. Cell, 146, 533–543.

27. Hamperl, S., Bocek, M.J., Saldivar, J.C., Swigut, T. and Cimprich, K.A. (2017) Transcription-Replication Conflict Orientation Modulates R-Loop Levels and Activates Distinct DNA Damage Responses. Cell, 170, 774–786 e719.

28. Zatreanu, D., Han, Z., Mitter, R., Tumini, E., Williams, H., Gregersen, L., Dirac-Svejstrup, A.B., Roma, S., Stewart, A., Aguilera, A. et al. (2019) Elongation Factor TFIIS Prevents Transcription Stress and R-Loop Accumulation to Maintain Genome Stability. Molecular cell, 76, 57–69 e59.

29. Lam, F.C., Kong, Y.W., Huang, Q., Vu Han, T.L., Maffa, A.D., Kasper, E.M. and Yaffe, M.B. (2020) BRD4 prevents the accumulation of R-loops and protects against transcription-replication collision events and DNA damage. Nat Commun, 11, 4083.

30. Shao, X., Joergensen, A.M., Howlett, N.G., Lisby, M. and Oestergaard, V.H. (2020) A distinct role for recombination repair factors in an early cellular response to transcription-replication conflicts. Nucleic Acids Res.

31. Sanchez, A., de Vivo, A., Tonzi, P., Kim, J., Huang, T.T. and Kee, Y. (2020) Transcription-replication conflicts as a source of common fragile site instability caused by BMI1-RNF2 deficiency. PLoS Genet, 16, e1008524.

32. Schroder, A.J., Pavlidis, P., Arimura, A., Capece, D. and Rothman, P.B. (2002) Cutting edge: STAT6 serves as a positive and negative regulator of gene expression in IL-4-stimulated B lymphocytes. J Immunol, 168, 996–1000.

33. Doyle, S.E., O’Connell, R., Vaidya, S.A., Chow, E.K., Yee, K. and Cheng, G. (2003) Toll-like receptor 3 mediates a more potent antiviral response than Toll-like receptor 4. J Immunol, 170, 3565–3571.

34. Manis, J.P., Dudley, D., Kaylor, L. and Alt, F.W. (2002) IgH class switch recombination to IgG1 in DNA-PKcs-deficient B cells. Immunity, 16, 607–617.

35. Sanz, L.A. and Chedin, F. (2019) High-resolution, strand-specific R-loop mapping via S9.6-based DNA-RNA immunoprecipitation and high-throughput sequencing. Nat Protoc, 14, 1734–1755.

36. Quinlan, A.R. and Hall, I.M. (2010) BEDTools: a flexible suite of utilities for comparing genomic features. Bioinformatics, 26, 841–842.

37. Van Rossum, G.a.D.J., Fred L. (1995). Centrum voor Wiskunde en Informatica Amsterdam.

38. Van Rossum, G.a.D., Fred L. (2009). CreateSpace, Scotts Valley, CA.

39. Virtanen, P., Gommers, R., Oliphant, T.E., Haberland, M., Reddy, T., Cournapeau, D., Burovski, E., Peterson, P., Weckesser, W., Bright, J. et al. (2020) SciPy 1.0: fundamental algorithms for scientific computing in Python. Nat Methods, 17, 261–272.

40. Harris, C.R., Millman, K.J., van der Walt, S.J., Gommers, R., Virtanen, P., Cournapeau, D., Wieser, E., Taylor, J., Berg, S., Smith, N.J. et al. (2020) Array programming with NumPy. Nature, 585, 357–362.

41. Hunter, J.D. (2007) Matplotlib: A 2D Graphics Environment. Computing in Science & Engineering, 9, 90–95.

42. McKinney, W. (2010) Data Structures for Statistical Computing in Python.

43. Oliphant, T.E. (2007) Python for Scientific Computing. Computing in Science & Engineering, 9, 10–20.

44. Karolchik, D., Hinrichs, A.S., Furey, T.S., Roskin, K.M., Sugnet, C.W., Haussler, D. and Kent, W.J. (2004) The UCSC Table Browser data retrieval tool. Nucleic Acids Res, 32, D493–496.

45. Kent, W.J., Zweig, A.S., Barber, G., Hinrichs, A.S. and Karolchik, D. (2010) BigWig and BigBed: enabling browsing of large distributed datasets. Bioinformatics, 26, 2204–2207.

46. Leinonen, R., Sugawara, H., Shumway, M. and International Nucleotide Sequence Database, C. (2011) The sequence read archive. Nucleic Acids Res, 39, D19–21.

47. Bolger, A.M., Lohse, M. and Usadel, B. (2014) Trimmomatic: a flexible trimmer for Illumina sequence data. Bioinformatics, 30, 2114–2120.

48. Mouse Genome Sequencing, C., Waterston, R.H., Lindblad-Toh, K., Birney, E., Rogers, J., Abril, J.F., Agarwal, P., Agarwala, R., Ainscough, R., Alexandersson, M. et al. (2002) Initial sequencing and comparative analysis of the mouse genome. Nature, 420, 520–562.

49. Langmead, B. and Salzberg, S.L. (2012) Fast gapped-read alignment with Bowtie 2. Nat Methods, 9, 357–359.

50. Li, H., Handsaker, B., Wysoker, A., Fennell, T., Ruan, J., Homer, N., Marth, G., Abecasis, G., Durbin, R. and Genome Project Data Processing, S. (2009) The Sequence Alignment/Map format and SAMtools. Bioinformatics, 25, 2078–2079.

51. Kent, W.J., Sugnet, C.W., Furey, T.S., Roskin, K.M., Pringle, T.H., Zahler, A.M. and Haussler, D. (2002) The human genome browser at UCSC. Genome Res, 12, 996–1006.

52. Feng, J., Liu, T., Qin, B., Zhang, Y. and Liu, X.S. (2012) Identifying ChIP-seq enrichment using MACS. Nat Protoc, 7, 1728–1740.

53. R Core Team. (2019). R Foundation for Statistical Computing, Vienna, Austria.

54. Stark, R. and Brown, G. (2011).

55. Ross-Innes, C.S., Stark, R., Teschendorff, A.E., Holmes, K.A., Ali, H.R., Dunning, M.J., Brown, G.D., Gojis, O., Ellis, I.O., Green, A.R. et al. (2012) Differential oestrogen receptor binding is associated with clinical outcome in breast cancer. Nature, 481, 389–393.

56. Heger, A., Webber, C., Goodson, M., Ponting, C.P. and Lunter, G. (2013) GAT: a simulation framework for testing the association of genomic intervals. Bioinformatics, 29, 2046–2048.

57. Law, M. and Shaw, D.R. (2018) Mouse Genome Informatics (MGI) Is the International Resource for Information on the Laboratory Mouse. Methods in molecular biology, 1757, 141–161.

58. Canela, A., Maman, Y., Jung, S., Wong, N., Callen, E., Day, A., Kieffer-Kwon, K.R., Pekowska, A., Zhang, H., Rao, S.S.P. et al. (2017) Genome Organization Drives Chromosome Fragility. Cell, 170, 507–521 e518.

59. Gardiner-Garden, M. and Frommer, M. (1987) CpG islands in vertebrate genomes. J Mol Biol, 196, 261–282.

60. Haeussler, M., Zweig, A.S., Tyner, C., Speir, M.L., Rosenbloom, K.R., Raney, B.J., Lee, C.M., Lee, B.T., Hinrichs, A.S., Gonzalez, J.N. et al. (2019) The UCSC Genome Browser database: 2019 update. Nucleic Acids Res, 47, D853–D858.

61. Barlow, J.H., Faryabi, R.B., Callen, E., Wong, N., Malhowski, A., Chen, H.T., Gutierrez-Cruz, G., Sun, H.W., McKinnon, P., Wright, G. et al. (2013) Identification of early replicating fragile sites that contribute to genome instability. Cell, 152, 620–632.

62. Le Tallec, B., Millot, G.A., Blin, M.E., Brison, O., Dutrillaux, B. and Debatisse, M. (2013) Common fragile site profiling in epithelial and erythroid cells reveals that most recurrent cancer deletions lie in fragile sites hosting large genes. Cell Rep, 4, 420–428.

63. Helmrich, A., Stout-Weider, K., Hermann, K., Schrock, E. and Heiden, T. (2006) Common fragile sites are conserved features of human and mouse chromosomes and relate to large active genes. Genome Res, 16, 1222–1230.

64. Shiraishi, T., Druck, T., Mimori, K., Flomenberg, J., Berk, L., Alder, H., Miller, W., Huebner, K. and Croce, C.M. (2001) Sequence conservation at human and mouse orthologous common fragile regions, FRA3B/FHIT and Fra14A2/Fhit. Proc Natl Acad Sci U S A, 98, 5722–5727.

65. Krummel, K.A., Denison, S.R., Calhoun, E., Phillips, L.A. and Smith, D.I. (2002) The common fragile site FRA16D and its associated gene WWOX are highly conserved in the mouse at Fra8E1. Genes Chromosomes Cancer, 34, 154–167.

66. Rozier, L., El-Achkar, E., Apiou, F. and Debatisse, M. (2004) Characterization of a conserved aphidicolin-sensitive common fragile site at human 4q22 and mouse 6C1: possible association with an inherited disease and cancer. Oncogene, 23, 6872–6880.

67. Helmrich, A., Stout-Weider, K., Matthaei, A., Hermann, K., Heiden, T. and Schrock, E. (2007) Identification of the human/mouse syntenic common fragile site FRA7K/Fra12C1--relation of FRA7K and other human common fragile sites on chromosome 7 to evolutionary breakpoints. Int J Cancer, 120, 48–54.

68. Ramirez, F., Dundar, F., Diehl, S., Gruning, B.A. and Manke, T. (2014) deepTools: a flexible platform for exploring deep-sequencing data. Nucleic Acids Res, 42, W187–191.

69. Tubbs, A., Sridharan, S., van Wietmarschen, N., Maman, Y., Callen, E., Stanlie, A., Wu, W., Wu, X., Day, A., Wong, N. et al. (2018) Dual Roles of Poly(dA:dT) Tracts in Replication Initiation and Fork Collapse. Cell.

70. Petryk, N., Kahli, M., d’Aubenton-Carafa, Y., Jaszczyszyn, Y., Shen, Y., Silvain, M., Thermes, C., Chen, C.L. and Hyrien, O. (2016) Replication landscape of the human genome. Nat Commun, 7, 10208.

71. Krupke, D.M., Begley, D.A., Sundberg, J.P., Richardson, J.E., Neuhauser, S.B. and Bult, C.J. (2017) The Mouse Tumor Biology Database: A Comprehensive Resource for Mouse Models of Human Cancer. Cancer Res, 77, e67–e70.

72. Newberg, J.Y., Mann, K.M., Mann, M.B., Jenkins, N.A. and Copeland, N.G. (2018) SBCDDB: Sleeping Beauty Cancer Driver Database for gene discovery in mouse models of human cancers. Nucleic Acids Res, 46, D1011–D1017.

73. Kieffer-Kwon, K.R., Nimura, K., Rao, S.S.P., Xu, J., Jung, S., Pekowska, A., Dose, M., Stevens, E., Mathe, E., Dong, P. et al. (2017) Myc Regulates Chromatin Decompaction and Nuclear Architecture during B Cell Activation. Molecular cell, 67, 566–578 e510.

74. Li, H. (2011) A statistical framework for SNP calling, mutation discovery, association mapping and population genetical parameter estimation from sequencing data. Bioinformatics, 27, 2987–2993.

75. Bergstrom, E.N., Huang, M.N., Mahto, U., Barnes, M., Stratton, M.R., Rozen, S.G. and Alexandrov, L.B. (2019) SigProfilerMatrixGenerator: a tool for visualizing and exploring patterns of small mutational events. BMC Genomics, 20, 685.

76. Kapusta, A., Kronenberg, Z., Lynch, V.J., Zhuo, X., Ramsay, L., Bourque, G., Yandell, M. and Feschotte, C. (2013) Transposable elements are major contributors to the origin, diversification, and regulation of vertebrate long noncoding RNAs. PLoS Genet, 9, e1003470.

77. Bedrat, A., Lacroix, L. and Mergny, J.L. (2016) Re-evaluation of G-quadruplex propensity with G4Hunter. Nucleic Acids Res, 44, 1746–1759.

78. Sernova, N.V. and Gelfand, M.S. (2008) Identification of replication origins in prokaryotic genomes. Brief Bioinform, 9, 376–391.

79. Heinz, S., Benner, C., Spann, N., Bertolino, E., Lin, Y.C., Laslo, P., Cheng, J.X., Murre, C., Singh, H. and Glass, C.K. (2010) Simple combinations of lineage-determining transcription factors prime cis-regulatory elements required for macrophage and B cell identities. Molecular cell, 38, 576–589.

80. Motenko, H., Neuhauser, S.B., O’Keefe, M. and Richardson, J.E. (2015) MouseMine: a new data warehouse for MGI. Mamm Genome, 26, 325–330.

81. Raudvere, U., Kolberg, L., Kuzmin, I., Arak, T., Adler, P., Peterson, H. and Vilo, J. (2019) g:Profiler: a web server for functional enrichment analysis and conversions of gene lists (2019 update). Nucleic Acids Res, 47, W191–W198.

82. Kuleshov, M.V., Jones, M.R., Rouillard, A.D., Fernandez, N.F., Duan, Q., Wang, Z., Koplev, S., Jenkins, S.L., Jagodnik, K.M., Lachmann, A. et al. (2016) Enrichr: a comprehensive gene set enrichment analysis web server 2016 update. Nucleic Acids Res, 44, W90–97.

83. Chen, E.Y., Tan, C.M., Kou, Y., Duan, Q., Wang, Z., Meirelles, G.V., Clark, N.R. and Ma’ayan, A. (2013) Enrichr: interactive and collaborative HTML5 gene list enrichment analysis tool. BMC Bioinformatics, 14, 128.

84. Mood, A.M. (1950) Introduction to the theory of statistics. 1st ed. McGraw-Hill, New York,.

85. Liu, B., Yi, J., Sv, A., Lan, X., Ma, Y., Huang, T.H., Leone, G. and Jin, V.X. (2013) QChIPat: a quantitative method to identify distinct binding patterns for two biological ChIP-seq samples in different experimental conditions. BMC Genomics, 14 Suppl 8, S3.

86. North, B.V., Curtis, D. and Sham, P.C. (2002) A note on the calculation of empirical P values from Monte Carlo procedures. Am J Hum Genet, 71, 439–441.

87. Rivals, I., Personnaz, L., Taing, L. and Potier, M.C. (2007) Enrichment or depletion of a GO category within a class of genes: which test? Bioinformatics, 23, 401–407.

88. Miura, M. and Chen, H. (2020) CUT&RUN detects distinct DNA footprints of RNA polymerase II near the transcription start sites. Chromosome Res.

89. Quinodoz, M., Gobet, C., Naef, F. and Gustafson, K.B. (2014) Characteristic bimodal profiles of RNA polymerase II at thousands of active mammalian promoters. Genome Biol, 15, R85.

90. Wissink, E.M., Vihervaara, A., Tippens, N.D. and Lis, J.T. (2019) Nascent RNA analyses: tracking transcription and its regulation. Nat Rev Genet.

91. Core, L.J., Waterfall, J.J., Gilchrist, D.A., Fargo, D.C., Kwak, H., Adelman, K. and Lis, J.T. (2012) Defining the status of RNA polymerase at promoters. Cell Rep, 2, 1025–1035.

92. Sanz, L.A., Hartono, S.R., Lim, Y.W., Steyaert, S., Rajpurkar, A., Ginno, P.A., Xu, X. and Chedin, F. (2016) Prevalent, Dynamic, and Conserved R-Loop Structures Associate with Specific Epigenomic Signatures in Mammals. Molecular cell, 63, 167–178.

93. Chen, Z., Li, S., Subramaniam, S., Shyy, J.Y. and Chien, S. (2017) Epigenetic Regulation: A New Frontier for Biomedical Engineers. Annu Rev Biomed Eng, 19, 195–219.

94. Sun, Z., Zhang, Y., Jia, J., Fang, Y., Tang, Y., Wu, H. and Fang, D. (2020) H3K36me3, message from chromatin to DNA damage repair. Cell Biosci, 10, 9.

95. Chen, Y.H., Keegan, S., Kahli, M., Tonzi, P., Fenyo, D., Huang, T.T. and Smith, D.J. (2019) Transcription shapes DNA replication initiation and termination in human cells. Nat Struct Mol Biol, 26, 67–77.

96. Desprat, R., Thierry-Mieg, D., Lailler, N., Lajugie, J., Schildkraut, C., Thierry-Mieg, J. and Bouhassira, E.E. (2009) Predictable dynamic program of timing of DNA replication in human cells. Genome Res, 19, 2288–2299.

97. Marchal, C., Sima, J. and Gilbert, D.M. (2019) Control of DNA replication timing in the 3D genome. Nat Rev Mol Cell Biol, 20, 721–737.

98. Macheret, M. and Halazonetis, T.D. (2019) Monitoring early S-phase origin firing and replication fork movement by sequencing nascent DNA from synchronized cells. Nat Protoc, 14, 51–67.

99. Toledo, L., Neelsen, K.J. and Lukas, J. (2017) Replication Catastrophe: When a Checkpoint Fails because of Exhaustion. Molecular cell, 66, 735–749.

100. Yu, C., Gan, H., Han, J., Zhou, Z.X., Jia, S., Chabes, A., Farrugia, G., Ordog, T. and Zhang, Z. (2014) Strand-specific analysis shows protein binding at replication forks and PCNA unloading from lagging strands when forks stall. Molecular cell, 56, 551–563.

101. Larsen, N.B., Liberti, S.E., Vogel, I., Jorgensen, S.W., Hickson, I.D. and Mankouri, H.W. (2017) Stalled replication forks generate a distinct mutational signature in yeast. Proc Natl Acad Sci U S A, 114, 9665–9670.

102. Franchitto, A. (2013) Genome instability at common fragile sites: searching for the cause of their instability. BioMed research international, 2013, 730714.

103. Bouwman, B.A.M. and Crosetto, N. (2018) Endogenous DNA Double-Strand Breaks during DNA Transactions: Emerging Insights and Methods for Genome-Wide Profiling. Genes (Basel), 9.

104. Vazquez, M.I., Catalan-Dibene, J. and Zlotnik, A. (2015) B cells responses and cytokine production are regulated by their immune microenvironment. Cytokine, 74, 318–326.

105. Canela, A., Sridharan, S., Sciascia, N., Tubbs, A., Meltzer, P., Sleckman, B.P. and Nussenzweig, A. (2016) DNA Breaks and End Resection Measured Genome-wide by End Sequencing. Molecular cell, 63, 898–911.

106. Casper, A.M., Nghiem, P., Arlt, M.F. and Glover, T.W. (2002) ATR regulates fragile site stability. Cell, 111, 779–789.

107. Morimoto, S., Tsuda, M., Bunch, H., Sasanuma, H., Austin, C. and Takeda, S. (2019) Type II DNA Topoisomerases Cause Spontaneous Double-Strand Breaks in Genomic DNA. Genes (Basel), 10.

108. Canela, A., Maman, Y., Huang, S.N., Wutz, G., Tang, W., Zagnoli-Vieira, G., Callen, E., Wong, N., Day, A., Peters, J.M. et al. (2019) Topoisomerase II-Induced Chromosome Breakage and Translocation Is Determined by Chromosome Architecture and Transcriptional Activity. Molecular cell, 75, 252–266 e258.

109. Dillon, L.W., Pierce, L.C., Ng, M.C. and Wang, Y.H. (2013) Role of DNA secondary structures in fragile site breakage along human chromosome 10. Hum Mol Genet, 22, 1443–1456.

110. Bird, A.P. (1980) DNA methylation and the frequency of CpG in animal DNA. Nucleic Acids Res, 8, 1499–1504.

111. Duncan, C.G., Kondilis-Mangum, H.D., Grimm, S.A., Bushel, P.R., Chrysovergis, K., Roberts, J.D., Tyson, F.L., Merrick, B.A. and Wade, P.A. (2018) Base-Resolution Analysis of DNA Methylation Patterns Downstream of Dnmt3a in Mouse Naive B Cells. G3 (Bethesda), 8, 805–813.

112. Ginno, P.A., Lim, Y.W., Lott, P.L., Korf, I. and Chedin, F. (2013) GC skew at the 5’ and 3’ ends of human genes links R-loop formation to epigenetic regulation and transcription termination. Genome Res, 23, 1590–1600.

113. Wang, Y., Yang, J., Wild, A.T., Wu, W.H., Shah, R., Danussi, C., Riggins, G.J., Kannan, K., Sulman, E.P., Chan, T.A. et al. (2019) G-quadruplex DNA drives genomic instability and represents a targetable molecular abnormality in ATRX-deficient malignant glioma. Nat Commun, 10, 943.

114. Tate, J.G., Bamford, S., Jubb, H.C., Sondka, Z., Beare, D.M., Bindal, N., Boutselakis, H., Cole, C.G., Creatore, C., Dawson, E. et al. (2019) COSMIC: the Catalogue Of Somatic Mutations In Cancer. Nucleic Acids Res, 47, D941–D947.

115. Le, H.P., Masuda, Y., Tsurimoto, T., Maki, S., Katayama, T., Furukohri, A. and Maki, H. (2015) Short CCG repeat in huntingtin gene is an obstacle for replicative DNA polymerases, potentially hampering progression of replication fork. Genes Cells, 20, 817–833.

116. Zhang, L. and Vijg, J. (2018) Somatic Mutagenesis in Mammals and Its Implications for Human Disease and Aging. Annu Rev Genet, 52, 397–419.

117. Hepburn, A.C., Steele, R.E., Veeratterapillay, R., Wilson, L., Kounatidou, E.E., Barnard, A., Berry, P., Cassidy, J.R., Moad, M., El-Sherif, A. et al. (2019) The induction of core pluripotency master regulators in cancers defines poor clinical outcomes and treatment resistance. Oncogene, 38, 4412–4424.

118. Feser, J. and Tyler, J. (2011) Chromatin structure as a mediator of aging. FEBS Lett, 585, 2041–2048.

119. Pannunzio, N.R. and Lieber, M.R. (2016) Dissecting the Roles of Divergent and Convergent Transcription in Chromosome Instability. Cell Rep, 14, 1025–1031.

120. Stolz, R., Sulthana, S., Hartono, S.R., Malig, M., Benham, C.J. and Chedin, F. (2019) Interplay between DNA sequence and negative superhelicity drives R-loop structures. Proc Natl Acad Sci U S A, 116, 6260–6269.

121. Ju, B.G., Lunyak, V.V., Perissi, V., Garcia-Bassets, I., Rose, D.W., Glass, C.K. and Rosenfeld, M.G. (2006) A topoisomerase IIbeta-mediated dsDNA break required for regulated transcription. Science, 312, 1798–1802.

122. Promonet, A., Padioleau, I., Liu, Y., Sanz, L., Biernacka, A., Schmitz, A.L., Skrzypczak, M., Sarrazin, A., Mettling, C., Rowicka, M. et al. (2020) Topoisomerase 1 prevents replication stress at R-loop-enriched transcription termination sites. Nat Commun, 11, 3940.

123. Rivera-Mulia, J.C. and Gilbert, D.M. (2016) Replication timing and transcriptional control: beyond cause and effect-part III. Curr Opin Cell Biol, 40, 168–178.

124. Wang, J., Rojas, P., Mao, J., Muste Sadurni, M., Garnier, O., Xiao, S., Higgs, M.R., Garcia, P. and Saponaro, M. (2021) Persistence of RNA transcription during DNA replication delays duplication of transcription start sites until G2/M. Cell Rep, 34, 108759.

125. Bhat, K.P. and Cortez, D. (2018) RPA and RAD51: fork reversal, fork protection, and genome stability. Nat Struct Mol Biol.

126. Biehs, R., Steinlage, M., Barton, O., Juhasz, S., Kunzel, J., Spies, J., Shibata, A., Jeggo, P.A. and Lobrich, M. (2017) DNA Double-Strand Break Resection Occurs during Non-homologous End Joining in G1 but Is Distinct from Resection during Homologous Recombination. Molecular cell, 65, 671–684 e675.

127. Seol, J.H., Shim, E.Y. and Lee, S.E. (2018) Microhomology-mediated end joining: Good, bad and ugly. Mutat Res, 809, 81–87.

128. Truong, L.N., Li, Y., Shi, L.Z., Hwang, P.Y., He, J., Wang, H., Razavian, N., Berns, M.W. and Wu, X. (2013) Microhomology-mediated End Joining and Homologous Recombination share the initial end resection step to repair DNA double-strand breaks in mammalian cells. Proc Natl Acad Sci U S A, 110, 7720–7725.

129. Vilette, D., Ehrlich, S.D. and Michel, B. (1996) Transcription-induced deletions in plasmid vectors: M13 DNA replication as a source of instability. Mol Gen Genet, 252, 398–403.

130. Viguera, E., Canceill, D. and Ehrlich, S.D. (2001) Replication slippage involves DNA polymerase pausing and dissociation. The EMBO journal, 20, 2587–2595.

131. Mao, Z., Bozzella, M., Seluanov, A. and Gorbunova, V. (2008) Comparison of nonhomologous end joining and homologous recombination in human cells. DNA Repair (Amst), 7, 1765–1771.

132. Merrikh, H., Machon, C., Grainger, W.H., Grossman, A.D. and Soultanas, P. (2011) Co-directional replication-transcription conflicts lead to replication restart. Nature, 470, 554–557.

133. Akamatsu, Y. and Kobayashi, T. (2015) The Human RNA Polymerase I Transcription Terminator Complex Acts as a Replication Fork Barrier That Coordinates the Progress of Replication with rRNA Transcription Activity. Molecular and cellular biology, 35, 1871–1881.

