## Supplementary figures for "Genomic patterns of transcription-replication interactions in mouse primary B cells"

**ST GERMAIN ET AL SUPPLEMENTARY DATA**

#### Supplementary Figure legends

**Supplementary Figure 1.** Transcription-Replication ImmunoPrecipitation on Nascent DNA followed by high throughput sequencing (TRIPn-Seq). **(A)** Representative UCSC genome browser tracks of RPM normalized reads showing results two methods of RNAPs5 ChIP-Seq. Yellow shading is showing additional signal in Method 2 not found in Method 1 and light blue shading is a TRI. **(B)** Properties of 357 TRIs found in an unbiased analysis. **(C)** Correlation plot of the analyzed regions between three TRIPn-Seq experiments and three controls. **(D)** Frequency histogram and table of gene length for TRI genes (green), all genes (red), and protein coding genes (black). P = $6.06 \times 10^{-40}$, Wilcoxon rank sum test. **(E)** Scatter plot and table displaying number of TRIs vs genes per chromosome.

**Supplementary Figure 2.** Characterization of transcriptional activity at TRIs. **(A)** Median Method 2 RNAP2s5 ChIP-Seq signal profiles scaled to full genes and centered near transcription termination sites TTSs for TRITSSs (green) and cTSSs (red). **(B)** Upper: Median EU-Seq signal profiles scaled to full genes for TRI genes (green) and cTSS genes (red). Lower: Median EU-Seq nascent RNA signal profiles centered near TSSs for plus strand TRITSSs (green), minus strand TRITSSs (light green), plus strand cTSSs (red), minus strand cTSSs (pink). **(C)** Upper: Median and GRO-Seq signal profiles centered near TSSs for plus strand TRIs (blue), minus strand TRIs (light blue), plus strand cTSSs (red), minus strand cTSSs (pink). Lower: Median GRO-Seq nascent RNA signal profiles for plus strand TRITSSs (green), minus strand TRITSSs (light green), plus strand cTSSs (red), minus strand cTSSs (pink). **(D)** DRIP-Seq signal heat maps centered near TSSs for TRIs, TRITSSs, and cTSSs. **(E)** Median DRIP-Seq signal profiles scaled to full genes (upper) and centered near TTSs (lower) for TRITSSs (green) and cTSSs (red). *All profile plot analyses are performed in mBCs. Shaded areas on the unfilled line plots are the standard error.

**Supplementary Figure 3.** Epigenetic landscape and chromatin modifier association at TRIs, TRITSSs and cTSS. **(A)** Median ChIP-Seq signal profiles of CTCF and GCN5. **(B)** Median ChIP-Seq signal profiles of histone acetylation marks H4K12Ac, H4K16Ac and H2bK20Ac as well as the histone deacetylase HDAC2. **(C)** Median ChIP-Seq signal profiles of H3K79me2, H3K27me2, H3K9me1 and H3K4me1/2/3. **(D)** Median ChIP-Seq signal profiles of H3K9me2 and H3K27me3. *All profile plot analyses are shown in RPKM, and performed in mBCs. Shaded areas on the unfilled line plots are the standard error.

**Supplementary Figure 4.** Replication characteristics at TRIs. **(A)** Median OK-Seq signal profile centered near TSSs of TRITSSs (green, n=583) and cTSSs (red, 6,558). **(B)** Median OK-Seq signal profiles scaled to full genes for TRI genes (green, n=583) and cTSS genes matched for transcriptional activity using EU-Seq (red, n=5,500). **(C)** Median EdC-Seq 10 mM HU signal profiles centered at TSSs for TRIs, TRITSSs, and cTSSs. **(D)** Median EdU-Seq 0 mM HU signal profiles centered at TSSs for TRIs, TRITSSs, and cTSSs. **(E)** Graphic representation (upper) and corresponding signal profiles (lower) of RPA accumulating more on the lagging strand of a right moving fork which is the plus strand (dark blue) and on the lagging strand of a left moving fork which is the minus strand (light blue). **(F)** Venn diagram showing the overlap between early replicating fragile sites (ERFSs) and TRIs in kilobase pairs (kbp), showing empirical p-value. **(G)** Venn diagram showing the overlap between common fragile sites (CFSs) and TRIs in kbp, showing empirical p-value.

**Supplementary Figure 5.** DNA breaks at TRIs. **(A)** Venn diagram showing the overlap between TRIs and END-Seq peaks, showing empirical p-value. **(B)** Median END-Seq +/- ATRi signal profiles centered near TSSs for TRIs (blue), TRITSSs (green), and cTSSs (red). **(C)** Median END-Seq signal profiles in WT and TOP2BKO cells +/- etoposide centered near TSSs for TRITSSs (green), cTSSs (red) and inactive TSSs (purple). **(D)** Median END-Seq 0 mM HU signal profiles centered near at forward strand TRITSSs (dark green), reverse strand TRITSSs (light green), forward strand cTSSs (red), and reverse strand cTSSs (dark red). Model of a break forming as a result of a stalled helicase (top) and a break forming as a result of a stalled DNA polymerase. *All profile plot analyses are performed in mBCs. Shaded areas on the unfilled line plots are the standard error.

**Supplementary Figure 6.** Sequences and mutations at TRIs. **(A)** Table and bar graph showing nucleotide content at TRIs, TRITSSs, and cTSSs. **(B)** Venn diagram showing the overlap between TRIs and CpG islands, showing empirical p-value. **(C)** Bar graph of single basepair substitutions (SBS) and short insertions/deletions (indels) at TRI genes and cTSS genes compared to all genes with RNAP2s5 signal, showing empirical p-value. **(D)** COSMIC SBS mutational signatures of TRIs, TRITSSs, and cTSSs in 1kb windows and TRI genes and cTSS genes. **(E)** COSMIC indel mutational signatures of TRIs, TRITSSs, and cTSSs in 1kb windows and TRI genes and cTSS genes. **(F)** Venn diagram showing the overlap between TRI genes and gene mutations in mouse tumors from the Mouse Tumor Biology (MTB) program, showing empirical p-value. **(G)** Venn diagram showing the overlap between TRIs and cancer drivers from the Sleeping Beauty Cancer Driver Database (SBCDD), showing empirical p-value.

**List of Supplementary tables (Provided as Excel document)**

**Supplementary Table S1.** Table of TRIs.

**Supplementary Table S2.** Table of gene type representation of TRIs.

**Supplementary Table S3.** Table of Motifs found at TRIs, TRITSSs, and cTSSs.

**Supplementary Table S4.** Table of variant types and percentages found at TRIs, TRITSSs, and cTSSs.

**Supplementary Table S5.** Table of tumor mutations and types at TRIs.

**Supplementary Table S6.** Table of MouseMine phenotypes enriched in TRIs.

**Supplementary Table S7.** Table of cancer associated gene enrichment in TRIs.

**Supplementary Table S8.** Table of gene expression and chromatin associated gene enrichment in TRIs.

**Supplementary Table S9.** Table of pluripotency of stem cells gene enrichment in TRIs.

### Supplementary Figure 1

**A**

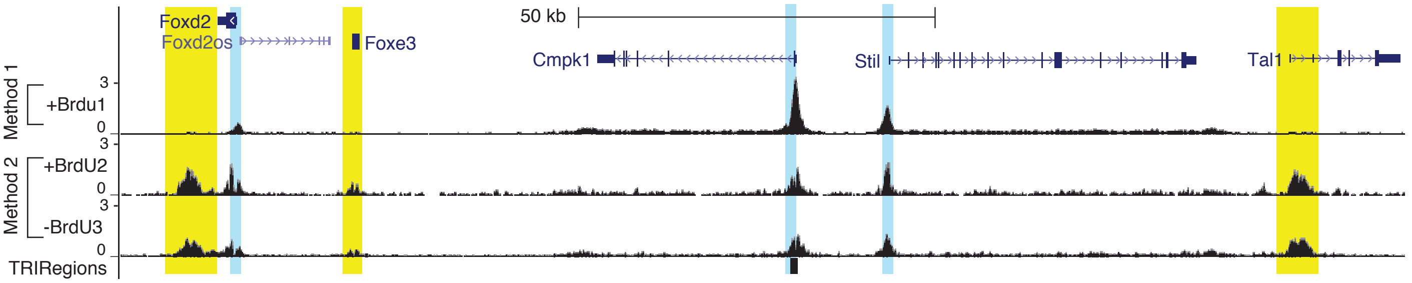


**B**

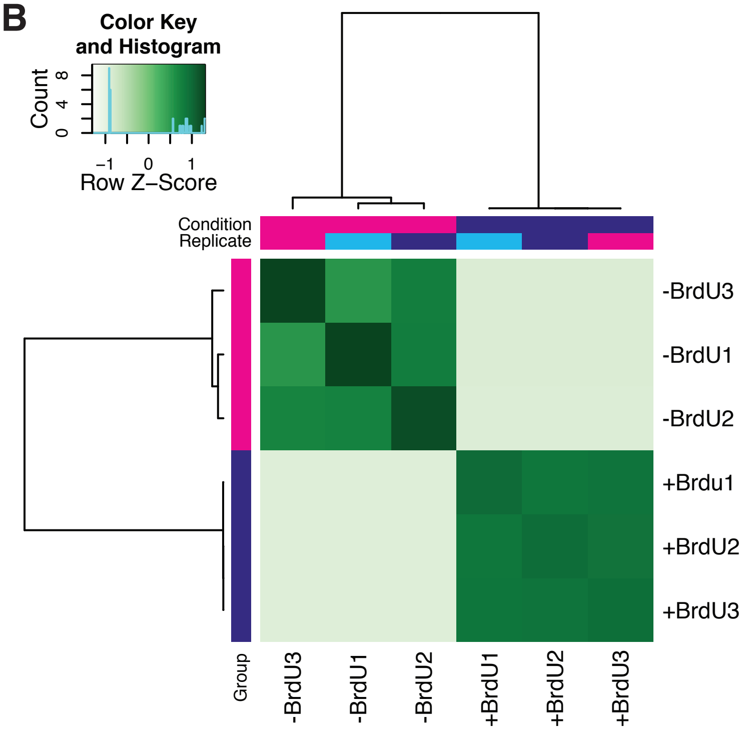


**C**

| TRI Properties | |
|---|---|
| # of input peaks | 76,154 |
| # with FDR < 0.05 | 357 |
| # overlapping TSS | 334 |
| Median (bp) | 528 |
| Mean (bp) | 588 |
| Min (bp) | 200 |
| Max (bp) | 1,676 |

**D**

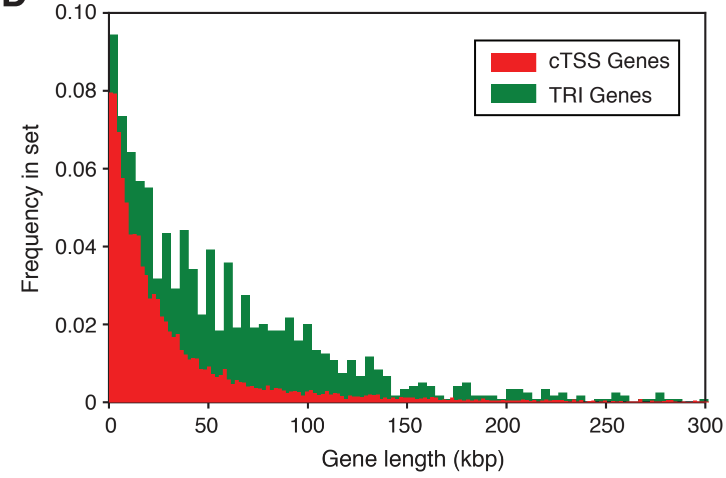


**E**

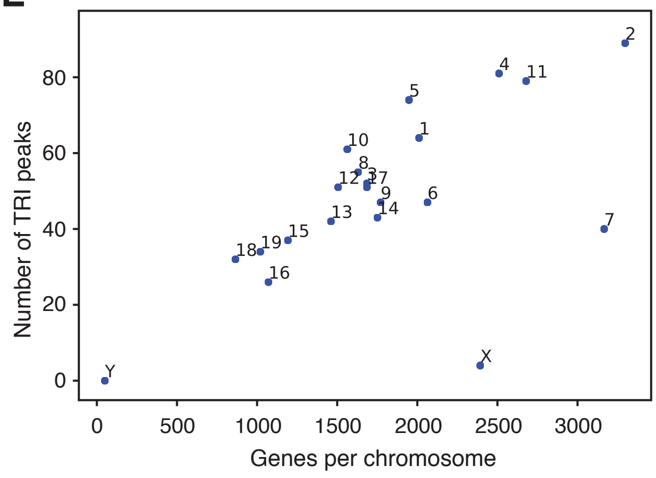

### Supplementary Figure 2

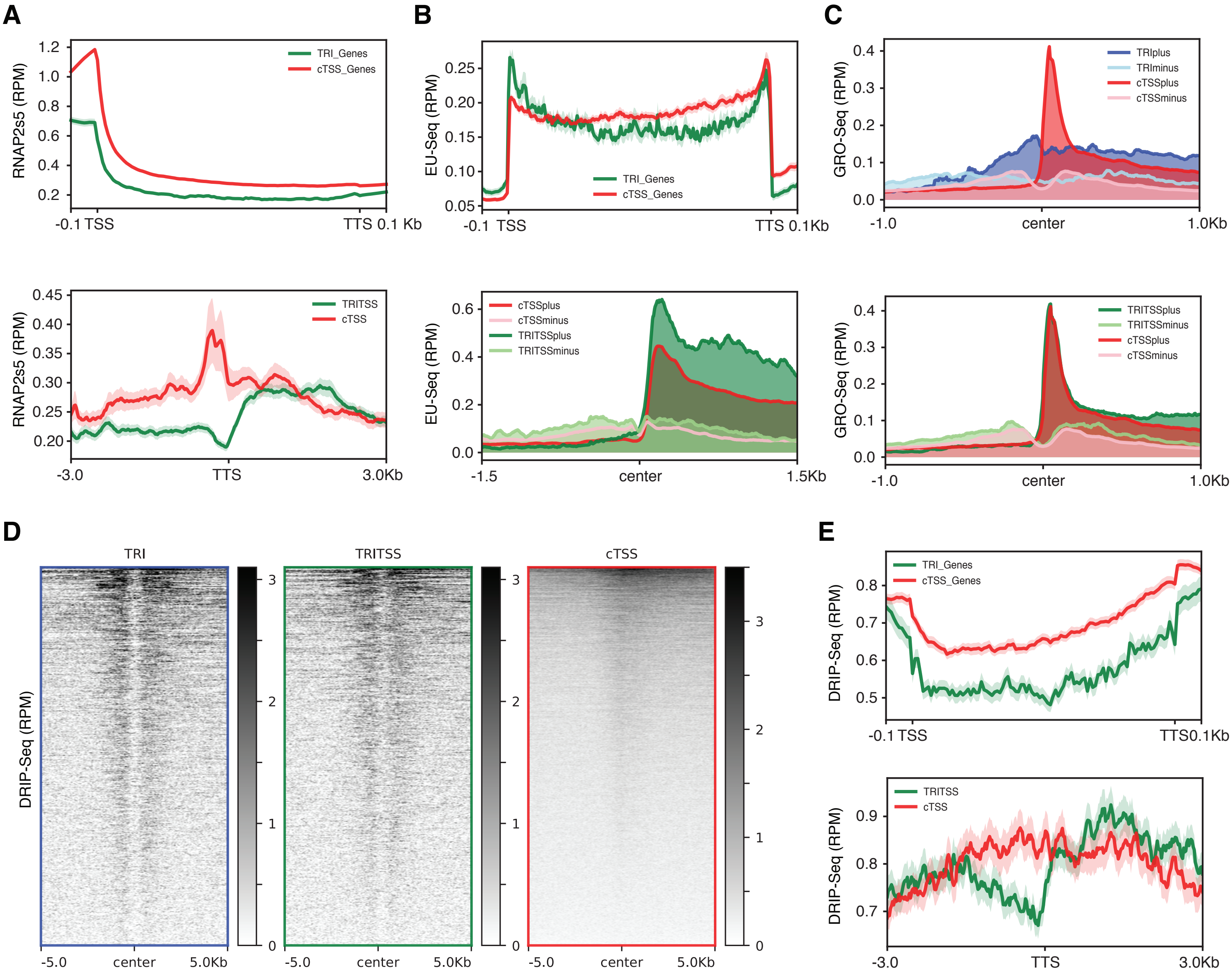

#### Supplementary figure 3

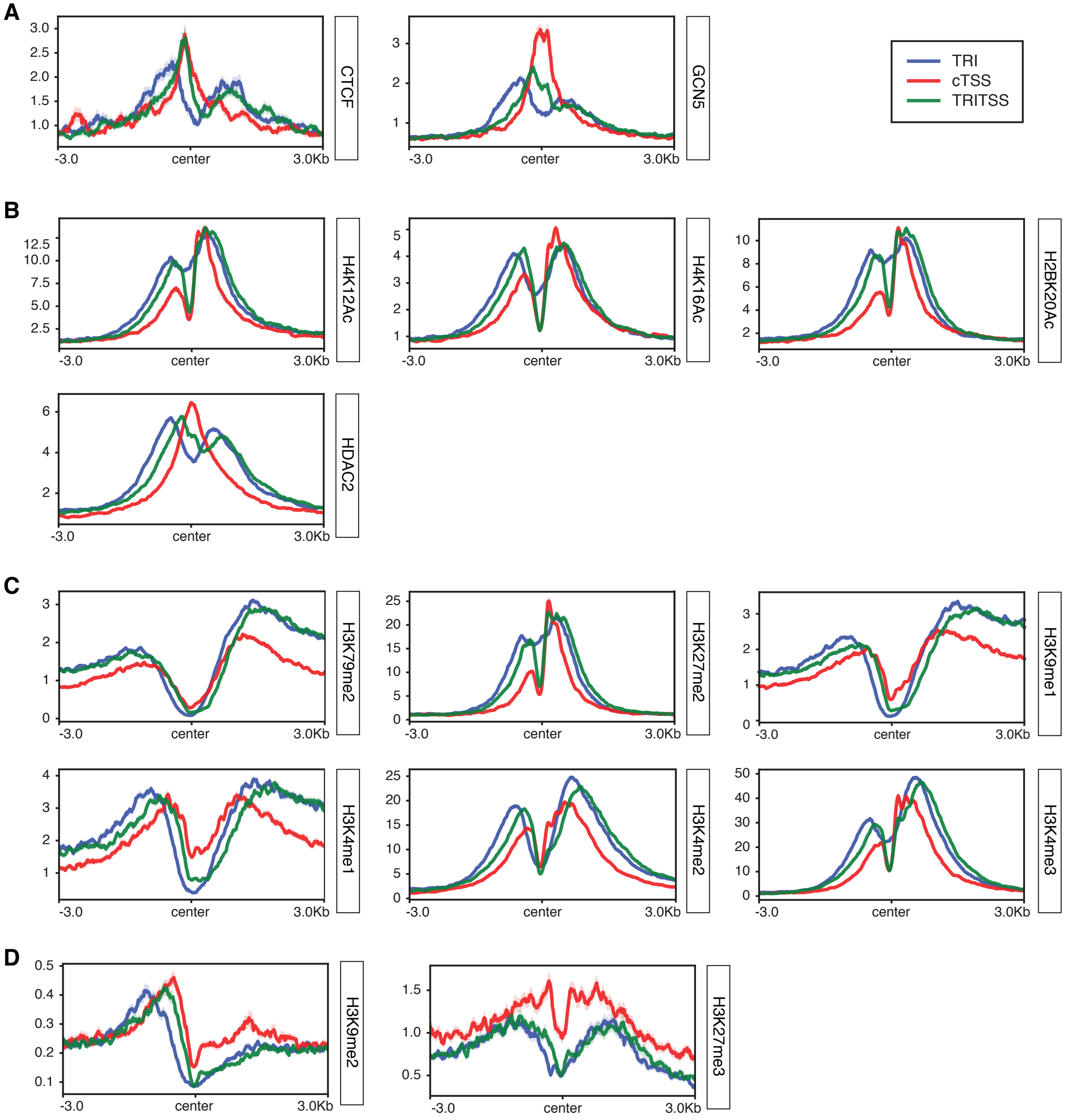

#### Supplementary Figure 4

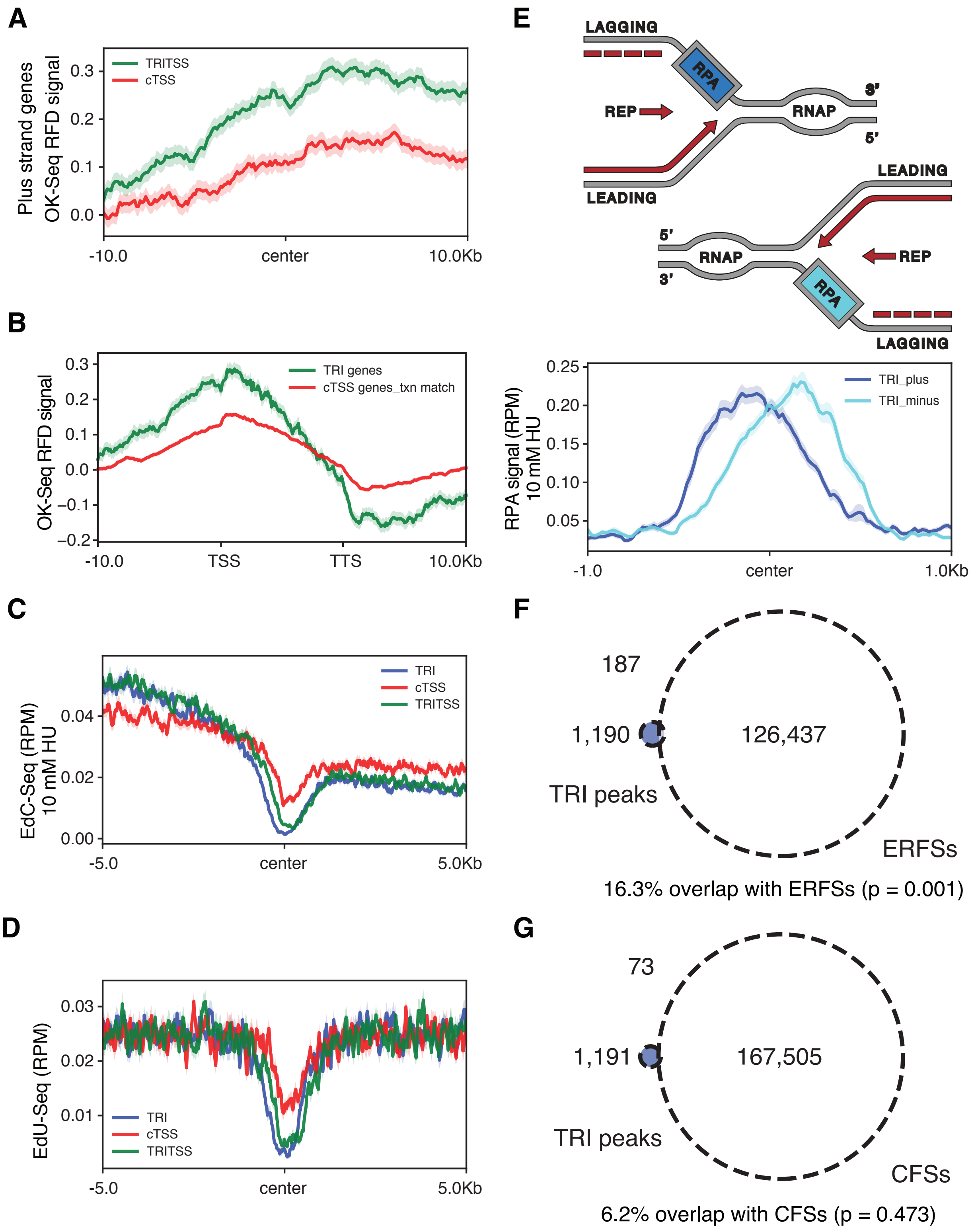

#### Supplementary Figure 5

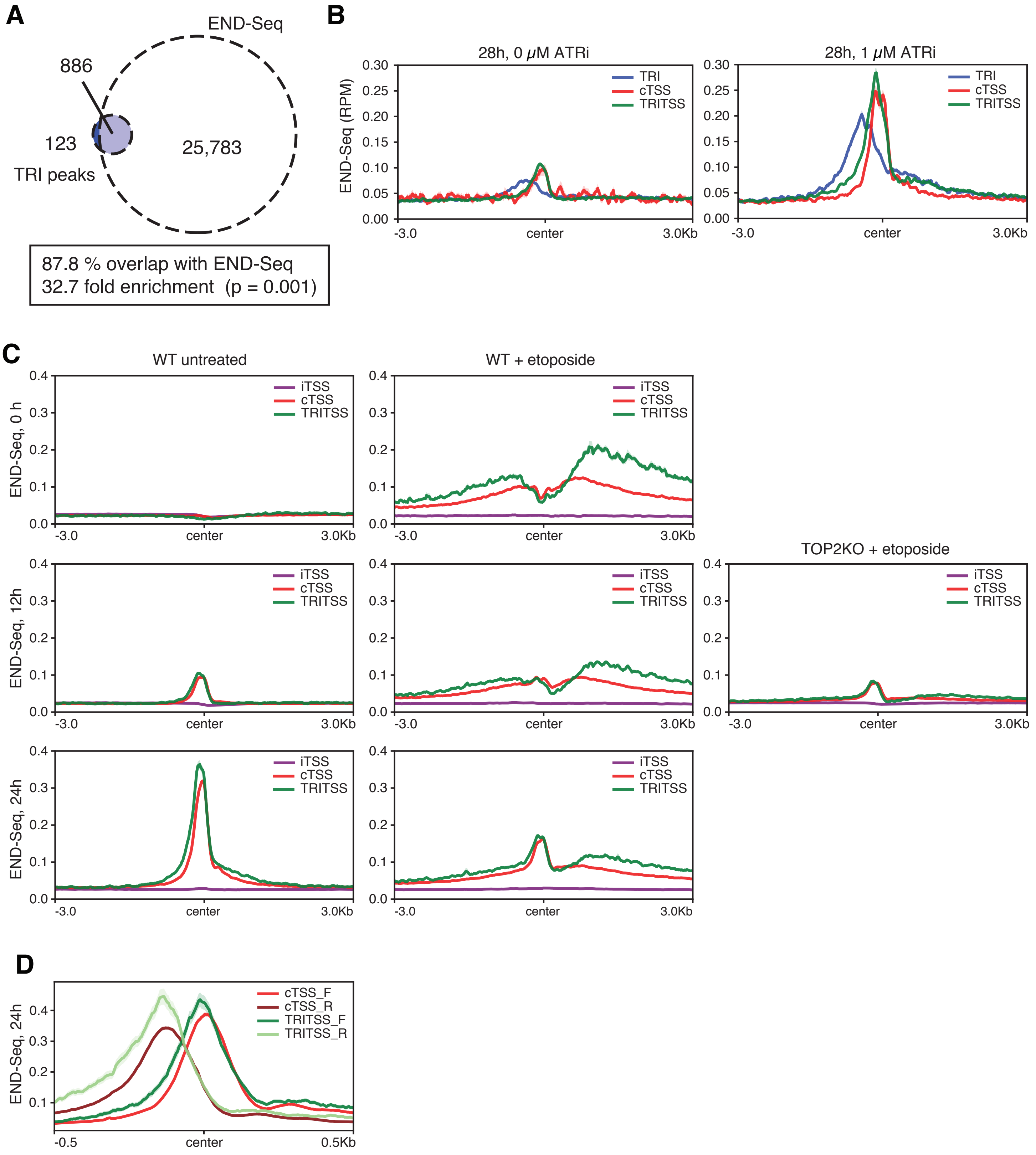

### Supplementary Figure 6

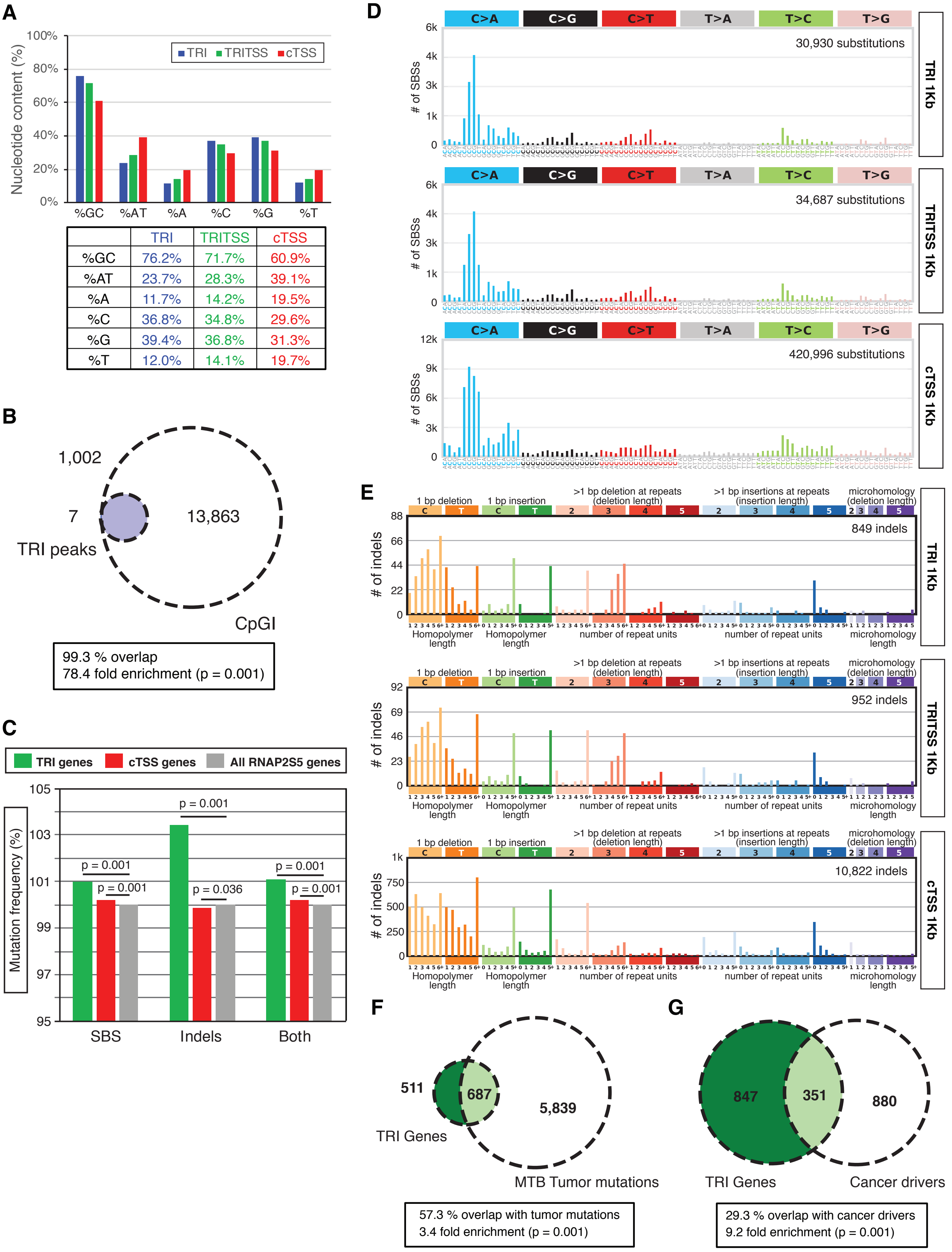


| | TRI | TRITSS | cTSS |
|---|---|---|---|
| %GC | 76.2% | 71.7% | 60.9% |
| %AT | 23.7% | 28.3% | 39.1% |
| %A | 11.7% | 14.2% | 19.5% |
| %C | 36.8% | 34.8% | 29.6% |
| %G | 39.4% | 36.8% | 31.3% |
| %T | 12.0% | 14.1% | 19.7% |
